# Spatial pharmaco-multiomics reveals drug distribution, metabolic niches, and spatially constrained resistance in medulloblastoma

**DOI:** 10.64898/2026.01.25.701559

**Authors:** Tuan Vo, Cedric S. Cui, Aitor Benedicto, Yibi Chen, Andrew Causer, Xiao Tan, Eun Ju Kim, Hani Vu, Albert Xiong, Mark Hodson, Yak Deng, Ana Maria Romero Jimenez, Ning Liu, Brett Hamilton, Thomas Robertson, Laura Genovesi, Brandon Wainwright, Jenny Fung, John Lee, Quan Nguyen

## Abstract

Single-cell and spatial transcriptomic studies have provided insights into the developmental origins and intratumoural heterogeneity of SHH medulloblastoma (SHH-MB) and suggested how targeted drugs such as CDK4/6 inhibitors remodel tumour ecosystems, yet the interplay between local drug exposure, metabolism, cell state, and drug resistance remains poorly understood. Here we developed a same-section spatial pharmaco-multiomics framework that integrates MALDI–MSI–based spatial metabolomics with Visium whole-transcriptome profiling and high-resolution Xenium imaging to map palbociclib distribution, metabolite landscapes, and transcriptional programs within the same histological contexts of an SHH-MB PDOX model and primary human tumours. Palbociclib-rich tumour bulk exhibited broad suppression of E2F-driven proliferation and a shift toward neuronal differentiation, corroborating and extending prior findings. In contrast, drug-poor tumour–brain interfaces and perivascular regions retained E2F-high proliferative states and were enriched for mesenchymal-like stromal cells and ECM-remodelling genes, indicating anatomically constrained reservoirs of tolerance. Spatial metabolomics linked these interface niches to ganglioside (GM2) and sphingomyelin enrichment, while differentiated, drug-exposed regions displayed phosphatidylcholine, phosphatidic-acid signatures consistent with neuronal maturation. Integrated pathway analysis further revealed a “mitochondrial tuning” program, with upregulation of histidine, folate/one-carbon, CoA, and lipoate metabolism with redox and oxidative-phosphorylation support. These signatures were specific to therapy-exposed border cells. Rare palbociclib-positive, E2F-high resistant spots additionally exhibited mitotic checkpoint and DNA-repair signatures, implying a drug-induced resistance axis independent of scarcity. Together, our study provides a generalisable same-section spatial pharmaco-multiomics pipeline and a spatially resolved model of CDK4/6 response, nominating interface-focused metabolic and cell-intrinsic vulnerabilities for combination therapy.

## Introduction

Medulloblastoma is the most common malignant paediatric brain tumour, characterised by profound heterogeneity and a complex tumour microenvironment (TME)^1,2^. The interplay between proliferative tumour cells, differentiating neuronal populations, and the extracellular matrix (ECM) is integral to maintaining tumour plasticity, enabling invasion, and fostering therapy resistance^2^. Among its subtypes, Sonic hedgehog medulloblastoma (SHH-MB) remains particularly challenging to treat due to spatial and cellular heterogeneity, which drives drug-tolerant states and relapse^3^. Resolving tumour heterogeneity and cell-state transitions at single-cell resolution is critical for elucidating resistance mechanisms and informing precision therapies. In addition to intrinsic heterogeneity, extrinsic factors such as tissue invasion and interactions with resident cells at tumour margins influence tumour progression, therapy response, and relapse^3,4^. For example, glial cells and glial-like tumour cells within the TME have been shown to promote SHH-MB growth through Hedgehog and other paracrine signalling mechanisms^3,5–7^. These observations underscore the need for approaches that capture both tumour-intrinsic and microenvironmental features *in situ*.

Palbociclib, a CDK4/6 inhibitor, has demonstrated efficacy in suppressing tumour proliferation and promoting differentiation in SHH-MB models^3^. Despite their promising results, approximately 80% of tumours relapse upon cessation of the Palbociclib treatment^8^, potentially due to residual proliferative cells in the interface regions^3^. Furthermore, patient derived xenograft models lacking an intact blood-brain barrier (BBB) suggest that drug distribution and microenvironmental interactions critically affect therapeutic outcomes^9^. These findings support the hypothesis that ECM-mediated interactions and vascular integrity enable transient drug-tolerant states, serving as a reservoir for relapse and potential drug resistance.

To address this, we developed a multimodal spatial-omics approach integrating spatial metabolomics and transcriptomics on the same tissue section. Spatial transcriptomics provides high-resolution mapping of gene expression within tissue contexts, revealing cellular states, cell-cell interactions, and signalling pathways that influence tumour behaviour and treatment response^3,10^. Spatial metabolomics complements this by profiling metabolites, lipids, and drug compounds, offering crucial insights into tumour metabolism and drug distribution^11,12^. Building on recent advances in spatial multi-omics^13–15^, our protocol enables concurrent mapping of transcriptomics and metabolomics profiles, allowing for direct association of drug distribution with adaptive responses in both tumour and resident cells. By investigating the spatial distribution of Palbociclib and its effects on the TME, this work provides insights into mechanisms driving therapy resistance and highlights potential strategies for overcoming therapeutic barriers in SHH-MB.

## Results

### Multimodal spatial omics framework for mapping CDK4/6 inhibitor response in MB-PDOX

To map how CDK4/6 inhibition reshapes tumour ecosystems *in situ*, we developed a same-section spatial pharmaco-multiomics workflow combining Matrix-Assisted Laser Desorption/Ionisation Mass Spectrometry Imaging (MALDI-MSI) spatial metabolomics with Visium^3^ whole-transcriptome and Xenium^10^ single-cell spatial transcriptomics (**Fig. 1a**). This approach enables high-resolution mapping of drug distribution, metabolite landscapes and transcriptional states across tumour and microenvironment. Species-resolved spatial transcriptomics enabled precise delineation of human tumour versus mouse stroma, consistent with pathologist-defined annotation, revealing distinct human, mouse and mixed spots at tumour-mouse interfaces (**Fig. 1b; Supplementary Figs. S1, S2**). Xenium further refined malignant versus host compartments at single-cell resolution (**Supplementary Fig. S3**). MALDI-MSI segmentation aligned with histology and species assignment, supporting robust integration across modalities (**Fig. 1b; Supplementary Figs. S1, S3**). Together, this multimodal framework enables high-resolution mapping of intratumoral heterogeneity, tumour-stroma interactions, and drug responses.

**Fig. 1:**
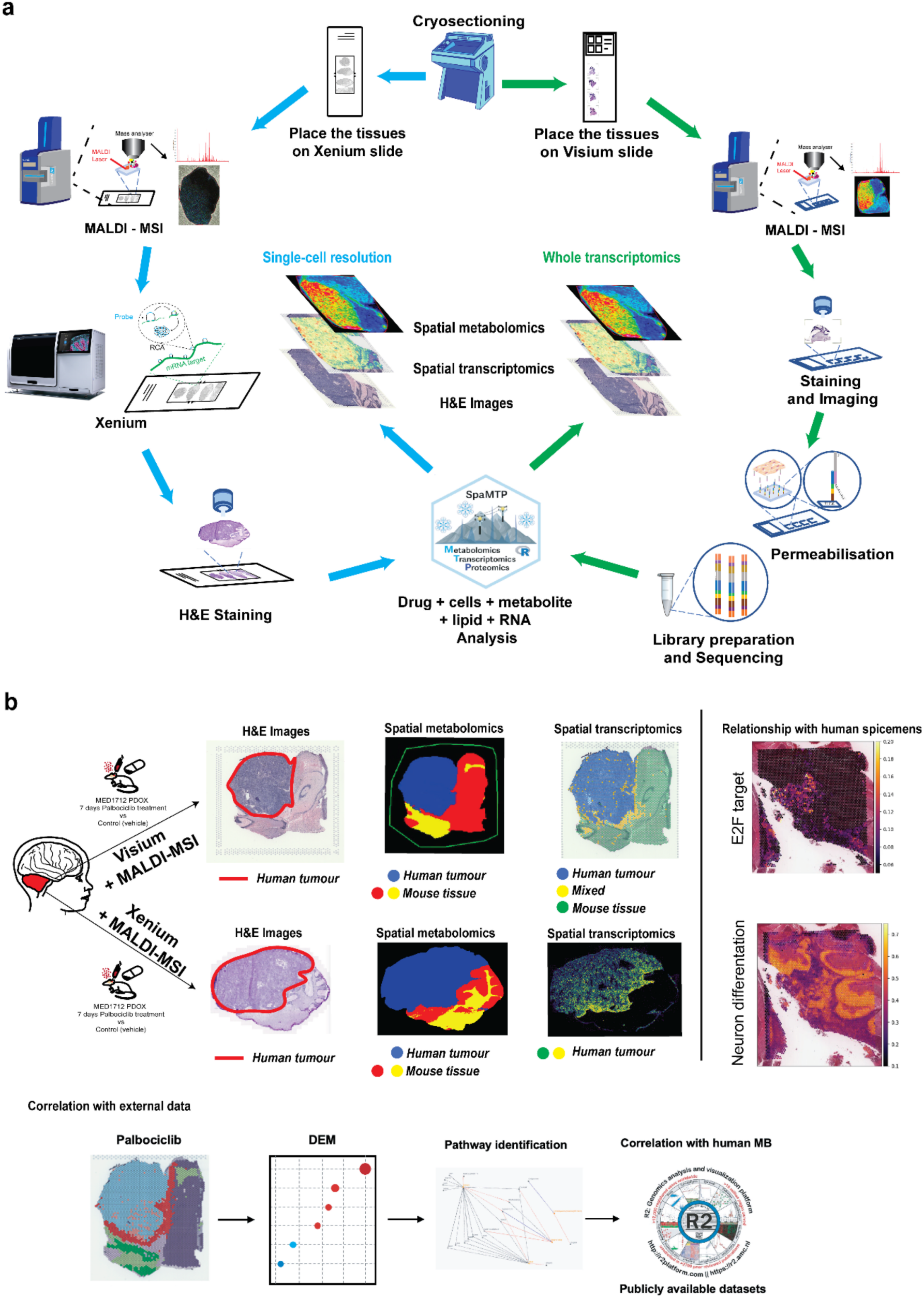
Workflow and experimental design for multimodal spatial omics in medulloblastoma PDOX models. **a** Medulloblastoma patient-derived orthotopic xenograft (PDOX) brains are cryo-sectioned and mounted either on Xenium slides for single-cell–resolution spatial transcriptomics or on Visium slides for whole-transcriptome spatial profiling. The sections on both slide types first undergo spatial metabolomic imaging by MALDI–MSI to map the distribution of drugs, metabolites and lipids (20 µm pixel size on Xenium slides and 50 µm on Visium slides). In the Xenium slide (left), MALDI–MSI is followed by high-resolution *in situ* transcriptomic profiling and subsequent H&E staining for histopathological assessment. In the Visium slide (right), sections are H&E-stained after MALDI–MSI and then processed through permeabilisation, library preparation and sequencing to obtain spot-resolved whole-transcriptome data. Joint analysis of MALDI–MSI and spatial transcriptomics using spaMTP package across both platforms enables integrated mapping of drug exposure, metabolic states and gene expression within heterogeneous tumour regions. **b** Spatial metabolomics and spatial transcriptomics are applied to medulloblastoma PDOX brains to distinguish human tumour from mouse host tissue and to characterise spatially resolved drug responses. The MED1712 PDOX model, derived from a paediatric medulloblastoma, is treated with the CDK4/6 inhibitor Palbociclib or vehicle -treated (control) for 7 days. Four SHH-MB mice (2 treated, 2 control) used for Visium, and four SHH-MB mice (2 treated, 2 control) used for Xenium. H&E staining shows tumour morphology, defining human tumour regions (red outline) and surrounding mouse brain. Spatial metabolomics segments tissue according to metabolite and lipid signatures, and spatial transcriptomics maps gene expression across the same sections, enabling concordant identification of human tumour and mouse compartments. Downstream analyses include differential metabolite and gene-expression profiling between treated and untreated tumour regions, pathway-level interrogation of drug-modulated programs. The PDOX spatial signatures are then compared with bulk and spatial transcriptomic datasets from human medulloblastoma to assess translational relevance.

### Spatial Palbociclib distribution shapes local tumour cell states

Palbociclib was predominantly detected in the tumour core and largely absent at tumour-mouse interfaces (**Fig. 2a, b; Supplementary Fig. S4**). Integrating these drug maps with spatial transcriptomics showed a strong inverse relationship between local drug exposure and proliferative programs: regions with high Palbociclib intensity displayed suppression of cell-cycle pathways, including *E2F* targets, and enrichment of neuronal differentiation and neurotransmitter transport signatures, while drug-poor interfaces retained proliferative activity (**Fig. 2c**; **Supplementary Fig. S5**). SHH-MB subtype mapping confirmed this spatial transition, with proliferative SHH-A signatures confined to interfaces and differentiated SHH-C signatures expanded in drug-rich tumour core (**Supplementary Fig. S6**).

**Fig. 2:**
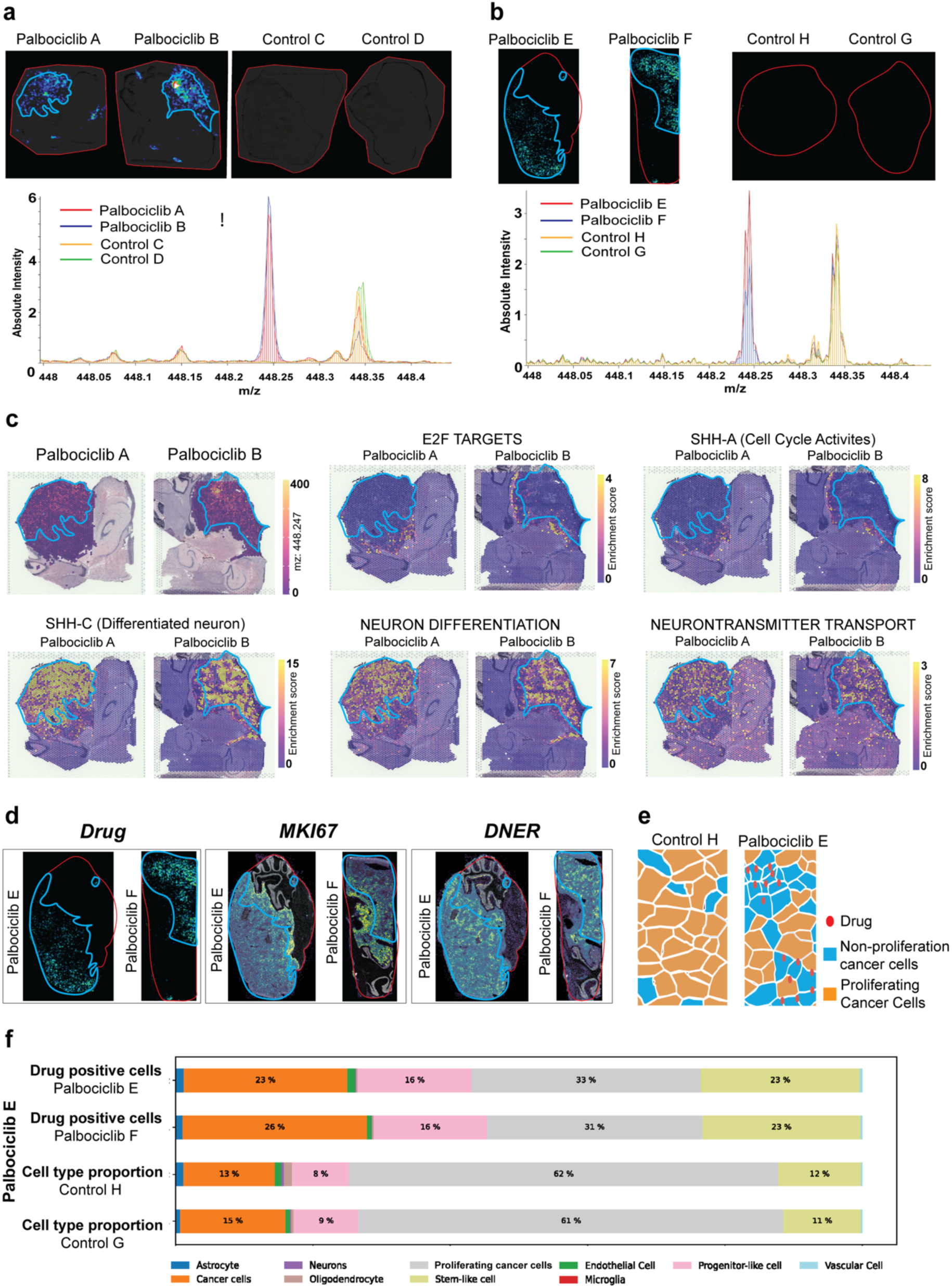
Spatial mapping of Palbociclib distribution and tumour cell states by integrated MALDI–MSI and spatial transcriptomics. **a** Drug detection by MALDI–MSI on Visium sections. The same tissue sections used for Visium spatial transcriptomics are subjected to MALDI–MSI to map Palbociclib. Ion images of the Palbociclib signal (m/z 448.25) are overlaid on brain sections with tissue borders (outlined in red), and intensity is rendered in blue. Control sections show no detectable Palbociclib signal. Representative mass spectra from tumour regions confirm a distinct Palbociclib peak in treated, but not control, samples. **b** Drug detection by MALDI–MSI on Xenium sections. MALDI–MSI performed on the same sections used for Xenium single-cell spatial transcriptomics similarly reveals robust Palbociclib signal in treated brains and absence of the m/z 448.25 peak in controls, demonstrating reproducible drug detection across both spatial transcriptomic platforms. **c** Overlay of drug distribution with Visium-derived tumour cell states. Drug-enriched regions (blue contours) defined from MALDI–MSI ion images are co-registered with Visium maps of E2F targets and SHH-A (cell cycle–active) modules, SHH-C (differentiated neuron) modules, neuron differentiation, and neurotransmitter transport gene-expression signatures. Palbociclib-exposed regions show **low** E2F targets and SHH-A scores and relatively **high** SHH-C, neuronal differentiation and neurotransmitter transport scores, whereas some highly proliferative peripheral regions remain drug-poor, consistent with microenvironmental constraints on drug penetration. **d** Overlay of drug distribution with proliferation and differentiation markers in Xenium data In Palbociclib-treated brains, drug-enriched areas (blue contours) on MALDI–MSI maps from the same sections correspond to reduced expression of the proliferation marker MKI67 and increased expression of the neuronal differentiation marker DNER at single-cell resolution, indicating local suppression of proliferative programs and enrichment of differentiated tumour cells in drug-exposed niches**. e** Cell type–resolved distribution of Palbociclib. A schematic tessellation map summarises Palbociclib-positive cells (red) across annotated cell types in control and treated tumours. Palbociclib signal is relatively depleted in proliferating tumour cells compared with non-proliferating tumour and non-malignant compartments, highlighting cell type– and location-dependent drug exposure. **f** Quantitative association between cell type and Palbociclib exposure. Stacked bar plots show, for each cell type, the fraction of cells classified as Palbociclib-positive based on spatial co-localisation with drug signal, together with overall cell-type composition in control and treated tumours. Palbociclib exposure is associated with a reduced proportion of proliferating tumour cells and a relative increase in differentiated and non-proliferating populations, consistent with effective targeting of cycling tumour cells in drug-accessible regions.

Single-cell Xenium data further showed inverse correlations between Palbociclib signal and proliferation marker *MKI67*, and positive correlations with differentiation markers such as *DNER* (**Fig. 2d**). Cell-type resolved analysis showed that Palbociclib-positive cells were relatively depleted among actively cycling tumour populations but more frequent in non-proliferative tumour cells and non-malignant compartments, including glial and vascular cells (**Fig. 2e**). Quantitative comparisons confirmed that treated tumours contained fewer proliferating tumour cells and a relative increase in differentiated and non-proliferating populations (**Fig. 2f**).

Together, these data indicate that Palbociclib remodels SHH-MB cell-states composition in a spatially structured manner: drug-accessible territories undergo cell-cycle arrest and neuronal differentiation, whereas drug-poor interfaces and perivascular niches remain proliferative, potentially sustaining treatment resistance.

### Spatial metabolomic niches define proliferative and differentiated tumour territories

Spatial metabolomics revealed regionally distinct tumour states aligned with histopathology and spatial transcriptomics **(Fig. 3a, b; Supplementary Fig. S2**). Unsupervised clustering of MALDI-MSI spectra identified seven metabolic clusters, including sphingomyelins/ubiquinol in the hippocampal pyramidal layer (Cluster 1, oxidative stress and mitochondrial activity)^16,17^, sphingomyelins/CDP-diacylglycerols in the cerebellar molecular layer (Cluster 3; lipid metabolism and synaptic activity)^18–20^, ceramides/phosphatidylethanolamines in brainstem structures (Cluster 4; lipid signalling and glial–neuronal interactions)^21–23^, and hexosylceramides/glucosylceramides in the cerebellar granule layer (Cluster 6; glycosphingolipid-dependent synaptic signalling)^24^. Specifically, Cluster 0 was enriched for ganglioside and sphingomyelins, marking proliferative territories^25–27^, and Cluster 2 was enriched for phosphatidylcholine and phosphatidic acids co-localising with Palbociclib exposure and neuronal differentiation signatures^28,29^. Treated samples were dominated by Cluster 2, whereas controls retained Cluster 0, indicating treatment-associated metabolic remodelling (**Fig. 3c)**. Differential analysis confirmed GM2 ganglioside enrichment in proliferative niches and Palbociclib enrichment in differentiated regions (**Fig. 3d**). Spatial mapping placed GM2-high territories at Palbociclib-poor tumour-mouse interfaces enriched for ECM-associated metabolites^30,31^, while tumour core displayed phospholipid profiles consistent with neuronal differentiation (**Fig. 3e, f**). Human SHH medulloblastoma specimens mirrored this organisation: spatial transcriptomics delineated zones of high *E2F* target expression versus neuronal differentiation scores, corresponding to proliferative and differentiated metabolic territories (**Fig. 3g, h**; **Supplementary Figs. S7 – S10**). Together, these findings demonstrate that Palbociclib remodels tumour metabolic states, shifting drug-accessible regions from proliferative, ganglioside-rich niches to differentiated territories, while drug-poor interfaces retain proliferative potential supported by ECM-associated metabolites.

**Fig. 3:**
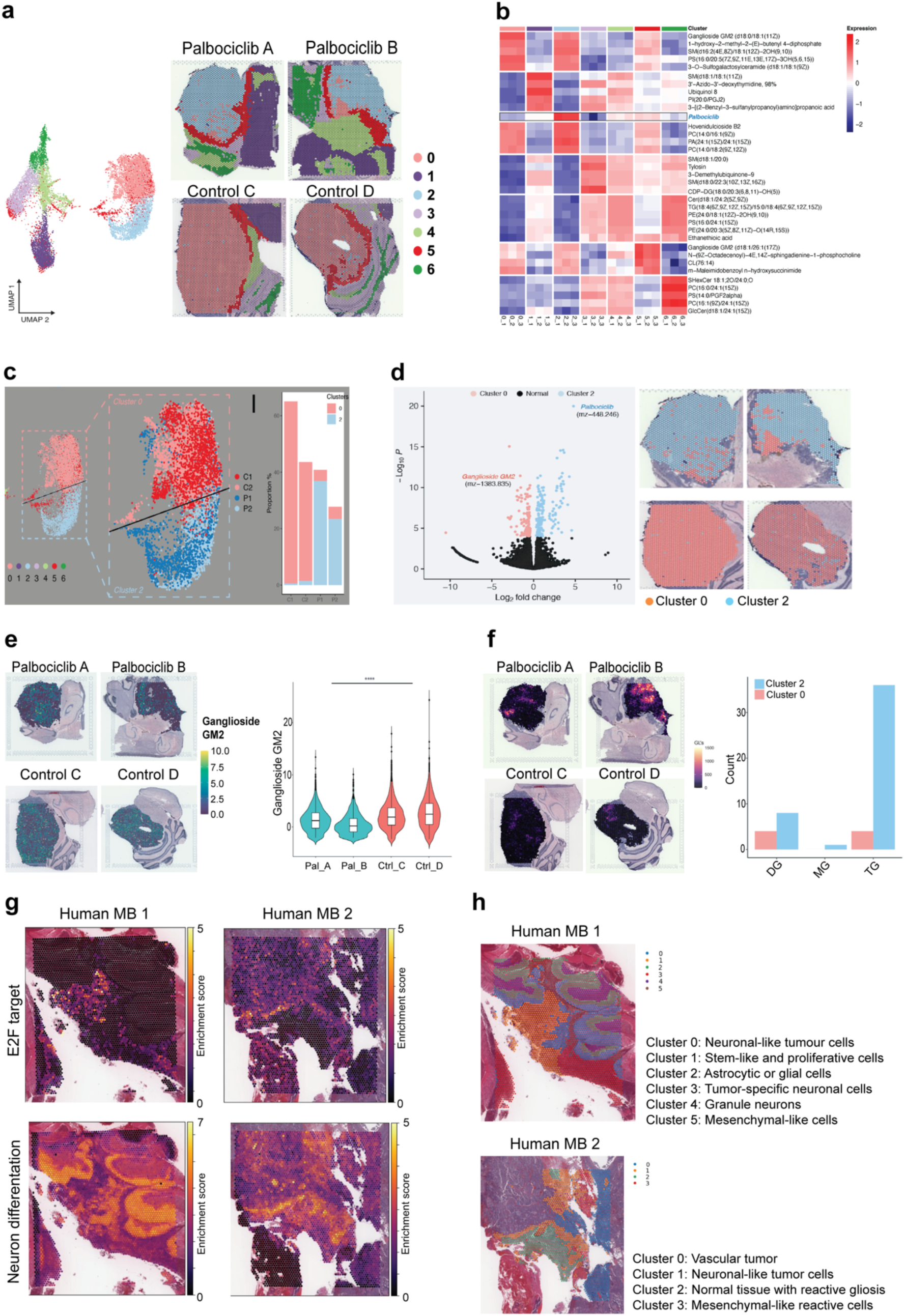
Spatial metabolomic states associated with proliferative programs in PDOX and human medulloblastoma. **a** Unsupervised clustering of MALDI–MSI data. UMAP embedding of MALDI–MSI spectra from Palbociclib-treated (Palbociclib A, Palbociclib B) and control (Control C, Control D) PDOX brains identifies seven metabolically distinct clusters. The corresponding spatial maps show the distribution of these clusters overlaid on H&E sections, revealing regionally segregated metabolic states in treated and untreated tumours. **b** Cluster-specific metabolite signatures. Hierarchical clustering heatmap summarising relative intensities of representative metabolites across the seven MALDI clusters. Clusters 0 and 2 are characterised by broadly elevated metabolite levels, with Cluster 2 distinguished by high Palbociclib signal compared with Cluster 0, consistent with distinct drug-exposed versus drug-poor metabolic states. **c** Sub-clustering of human tumour regions MALDI–MSI data restricted to human tumour areas (Clusters 0 and 2) from four PDOX samples are visualised in UMAP space. Treated samples (Palbociclib A, Palbociclib B) are enriched for non-proliferative Cluster 2, whereas control samples (Control C, Control D) are dominated by proliferative Cluster 0. Bar plots quantify the proportion of tumour pixels assigned to each cluster, highlighting treatment-associated remodelling of tumour metabolic states. **d** Metabolite differences between proliferative and non-proliferative states. Volcano plot of differential metabolite abundance between Cluster 0 (proliferative) and Cluster 2 (non-proliferative) tumour regions. Ganglioside GM2 (m/z 1,383.835) is significantly enriched in proliferative Cluster 0, whereas Palbociclib (m/z 448.246) is enriched in non-proliferative Cluster 2. Spatial maps illustrate the complementary distribution of these two clusters within representative tumours. **e** Ganglioside GM2–rich interfaces and extracellular matrix metabolites. Spatial maps show that ganglioside GM2–high regions are concentrated at tumour–brain interfaces and invasive fronts. Violin plots of selected extracellular matrix–associated metabolites demonstrate higher abundance in GM2-high proliferative regions compared with GM2-low non-proliferative regions, implicating a ganglioside–ECM niche in supporting tumour proliferation. **f** Glycolipid-enriched regions. Spatial maps and quantification of additional glycolipids reveal their preferential localisation to proliferative Cluster 0 regions, with reduced abundance in non-proliferative Cluster 2, further linking glycolipid metabolism to proliferative tumour niches. **g** Mapping of proliferative and differentiated cell states in human SHH medulloblastoma. In two primary SHH medulloblastoma specimens (Human MB 1 and Human MB 2), gene-set scores for E2F targets (top row) and neuron differentiation (bottom row) derived from spatial transcriptomics delineate discrete zones of high proliferation versus neuronal differentiation, mirroring the proliferative and non-proliferative metabolic territories observed in PDOX models. **h** Spatial heterogeneity of tumour and microenvironmental cell states in human SHH medulloblastoma. Clustering of spatial transcriptomic spots in the same specimens identifies multiple transcriptional states, including vascular tumour cells, cycling tumour cells, differentiated tumour cells and diverse non-malignant microenvironmental populations. Spatial maps of these clusters illustrate complex intratumoural heterogeneity and the interdigitation of proliferative, differentiated and stromal niches within the human medulloblastoma tumour microenvironment.

### Integrated transcriptomic–metabolomic pathway analysis identifies coordinated sphingolipid and differentiation programs

Joint pathway analysis across spatial transcriptomics and metabolomics revealed coordinated treatment-induced shifts coupling metabolism and cell-state (**Fig. 4a – c**). Among these, sphingolipid metabolism emerged as a central axis linking metabolic reprogramming to neuronal differentiation: combined metabolite-gene networks showed loss of pro-proliferative sphingolipid species and enrichment of differentiation-associated lipids in drug-exposed territories (**Fig. 4d**), consistent with membrane remodelling required for neuronal maturation. Spatial mapping confirmed contraction of proliferation-associated *HMGB2* and expansion of neuronal markers *NNAT* and *TUBA1B* within regions undergoing sphingolipid remodelling (**Fig. 4e**). Three-dimensional co-localisation of metabolites such as L-arginine with neuronal genes including *KIF5C* delineated contiguous “differentiation niches” in the tumour core (**Fig. 4f**). Together, these results demonstrate that Palbociclib induces a coordinated metabolic-transcriptional shift in drug-accessible tumour territories, coupling sphingolipid remodelling with robust neuronal differentiation programs to establish spatially organised differentiation niches within the tumour microenvironment.

**Fig. 4:**
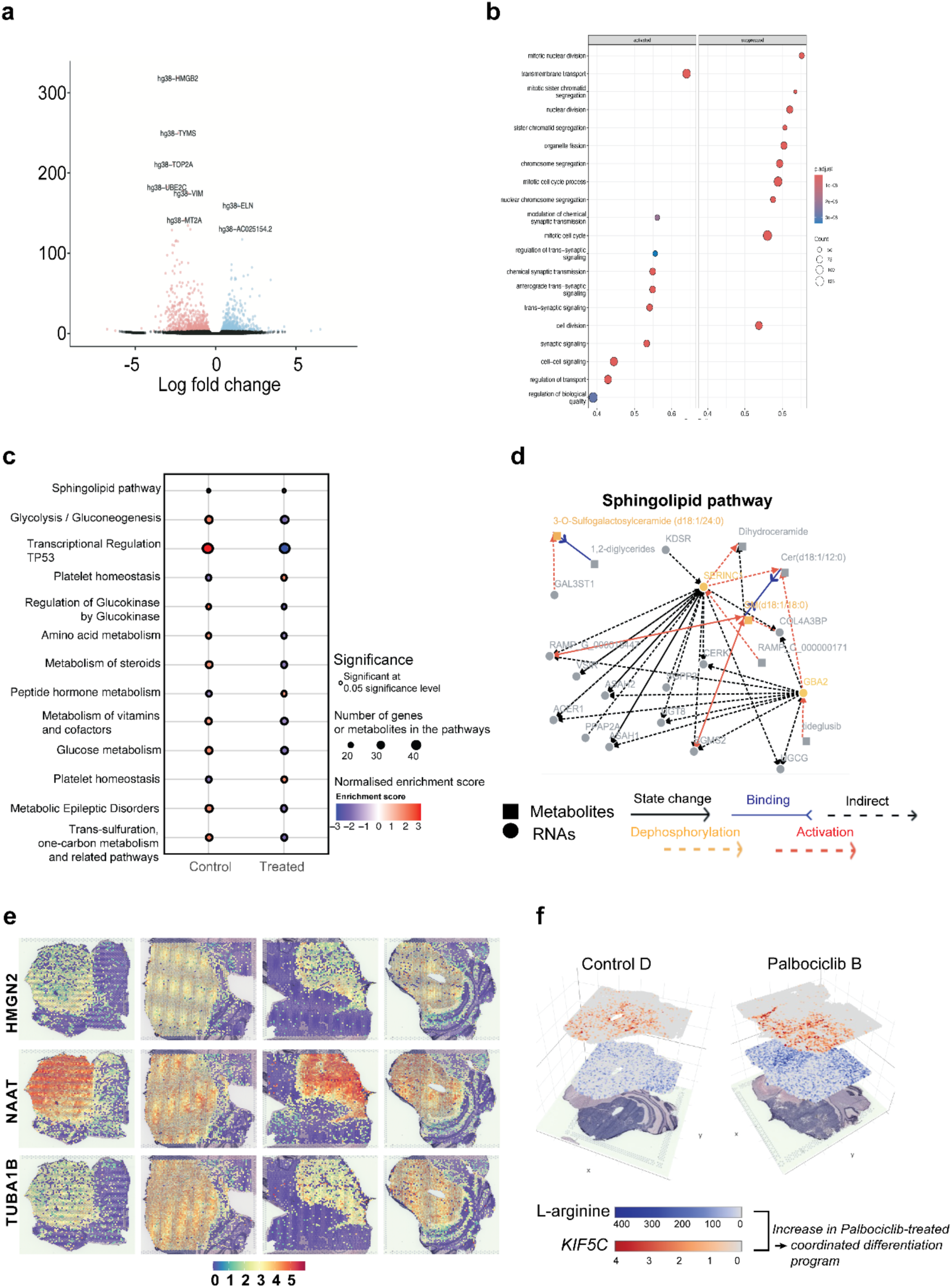
Integrated transcriptomic–metabolomic pathway analysis reveals a Palbociclib-induced differentiation programme. **a** Differential gene expression. Volcano plot showing differential gene expression between Palbociclib-treated and control tumour regions in medulloblastoma PDOX brains. Red and blue points denote significantly up- and downregulated genes, respectively (FDR-adjusted threshold), highlighting extensive transcriptional remodelling after treatment. **b** Pathway enrichment analysis. Dot plot of gene-set enrichment analysis for differentially expressed genes. Selected pathways involved in sphingolipid metabolism, glycolysis/gluconeogenesis, amino-acid metabolism, transcriptional regulation and platelet/vascular homeostasis are significantly enriched, with dot size indicating the number of altered genes and colour reflecting normalised enrichment scores. **c** Clustered pathway enrichment map. Comparison of pathway-level enrichment scores between control and Palbociclib-treated tumour regions. The sphingolipid pathway, glycolysis/gluconeogenesis and several metabolic and signalling pathways show coordinated shifts with treatment, indicating that Palbociclib reshapes both metabolic and regulatory programs. **d** Representative integrated pathway network (sphingolipid). Network representation of metabolites (circles) and RNAs (squares) in the sphingolipid pathway, integrating MALDI–MSI and spatial transcriptomic data. Edges denote known biochemical relationships, with colour and line style indicating activation, dephosphorylation, binding or indirect interactions. State changes (control versus treated) highlight a coherent reconfiguration of sphingolipid metabolism in drug-exposed tumour regions. **e** Spatial feature plots of selected marker genes. Visium spatial maps of HMGB2, NNAT and TUBA1B expression across PDOX sections from control and Palbociclib-treated brains. HMGB2 marks proliferative tumour zones that are reduced after treatment, whereas NNAT and TUBA1B, associated with neuronal differentiation, are increased in drug-exposed regions, consistent with a shift towards a differentiated state. **f** Three-dimensional overlay of gene and metabolite distributions. 3D renderings of a control tumour (Control D) and a Palbociclib-treated tumour (Palbociclib B) showing spatial co-localisation of the metabolite L-arginine (purple scale) and the neuronal differentiation gene KIF5C (red scale). Palbociclib-treated tumours display expanded regions with concordant elevation of L-arginine and KIF5C, illustrating a coordinated differentiation programme coupling metabolic and transcriptional changes in drug-responsive niches.

### Human-like tumour–brain interfaces form a distinct metabolic niche

Species-resolved spatial analyses revealed two metabolically distinct compartments at the tumour-mouse boundary (**Fig. 5a – f**). The human-like interface (tumour-dominant) displayed a cancer-linked metabolic program enriched for glycosphingolipid, fatty acid, nucleotide and arachidonic acid pathways, consistent with membrane biogenesis, DNA synthesis and pro-tumorigenic signalling at the invasive front. In contrast, the mouse-like interface (brain-dominant) was characterised by homeostatic pathways supporting neural integrity, including phosphatidylcholine and phosphatidylethanolamine metabolism, compatible with sustained mitochondrial function and neuroprotection in reactive astrocytes. Differential metabolite mapping highlighted species that discriminate these compartments: GM2 ganglioside, previously linked to proliferative and ECM-rich states, was selectively enriched in the human-like interface, whereas N-acetyl-L-aspartic acid, associated with neuronal integrity, localised to the mouse-like interface (**Fig. 5g – i**). Together, these findings demonstrate that the tumour-mouse boundary is partitioned into a cancer-metabolic niche and an adjacent brain-homeostatic niche, each defined by distinct lipid and metabolite profiles uncovered by our integrated Visium–MALDI platform.

**Fig. 5:**
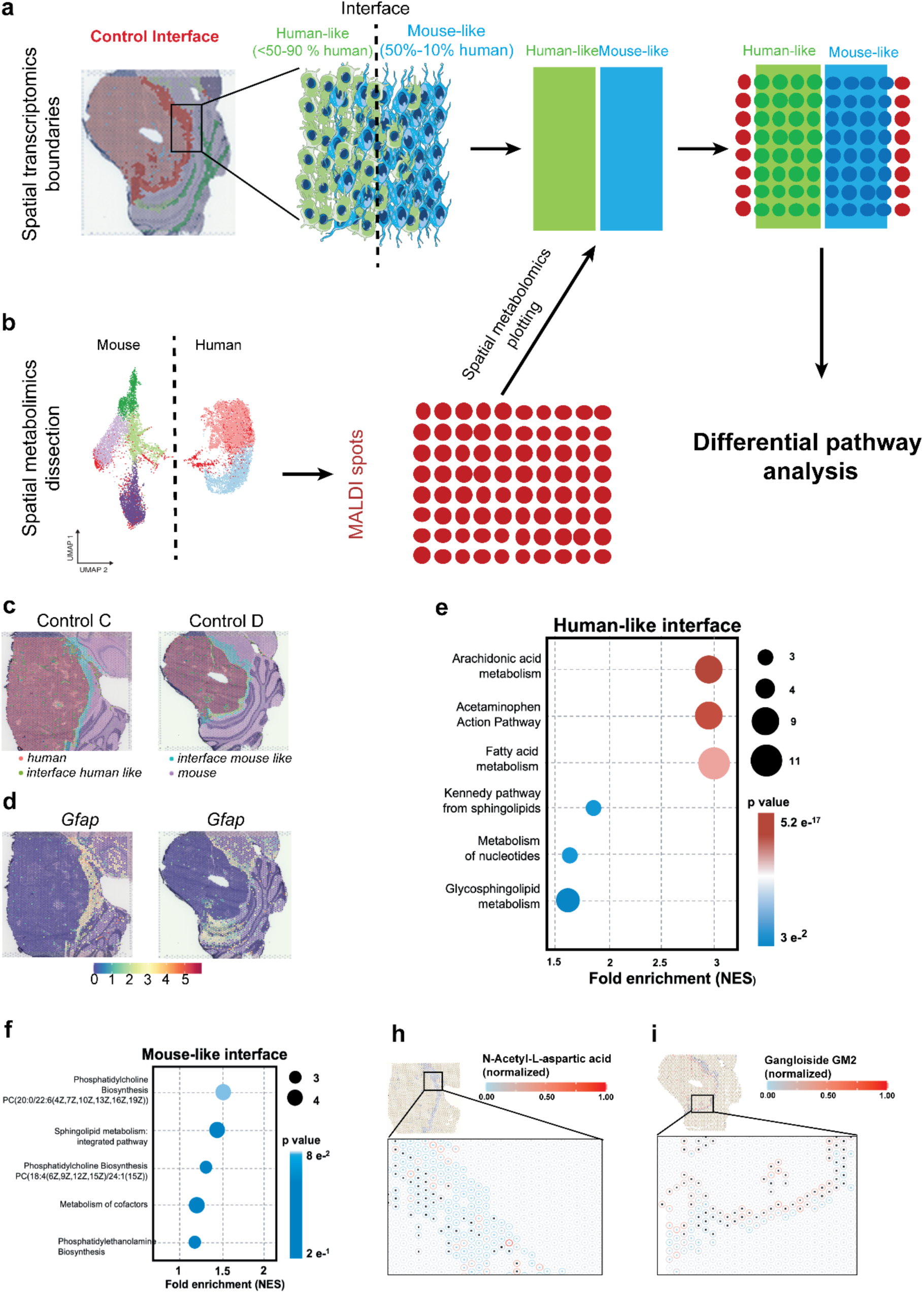
Spatial metabolomic profiling of tumour–brain interfaces in medulloblastoma PDOX. **a**. Defining human-like and mouse-like interfaces by spatial transcriptomics. Spatial transcriptomic spots are first classified as human tumour or mouse brain based on species-specific transcript abundance, and tumour boundaries are delineated. Spots with predominantly human tumour signal form the human-like compartment, those with predominantly mouse signal form the mouse-like compartment, and a mixed band of spots at the tumour margin defines the tumour–brain interface. These three regions are used as anatomical priors for downstream metabolomic and pathway analyses. **b** Mapping metabolites onto tumour-brain interfaces MALDI–MSI spectra acquired from the same sections are embedded in low-dimensional space to separate mouse- and human-associated metabolic profiles. Co-registration with spatial transcriptomic boundaries assigns each MALDI pixel to human-like, mouse-like or interface regions, enabling interface-specific metabolite and pathway analysis. **c** Spatial distribution of tumour–brain interfaces. Representative control brains (Control C and Control D) show the organisation of human tumour core, human-like interface and mouse-like interface regions. Human tumour cells extend into a narrow human-like interface adjacent to surrounding mouse brain tissue. **d** *Gfap* enrichment at the interface. Spatial maps of *Gfap* expression illustrate strong enrichment in the interface regions, consistent with reactive astrocytosis at the tumour–brain border and validating the interface definition derived from mixed human–mouse transcriptional content. **e** Cancer-linked metabolic pathways in the human-like interface. Gene set–style enrichment analysis of MALDI-derived metabolites assigned to the human-like interface identifies pathways associated with cancer metabolism, including arachidonic acid metabolism, fatty-acid metabolism, nucleotide metabolism and glycosphingolipid. Dot size indicates the number of altered metabolites per pathway and colour encodes the normalised enrichment score. **f** Brain-specific metabolic pathways in the mouse-like interface. An analogous enrichment analysis for metabolites in the mouse-like interface reveals phosphatidylcholine, Sphingolipid metabolism, phosphatidylethanolamine, reflecting the metabolic signature of surrounding normal brain tissue. **h,i** Representative metabolites distinguishing mouse-like and human-like interfaces. Spatial maps and corresponding quantification for exemplar metabolites show preferential accumulation in mouse-like (h) or human-like (i) interface regions, illustrating how distinct metabolic niches segregate across the tumour–brain border. Spots in human-like interface regions are marked with dark squares in the centre for both plots.

### Mitochondrial tuning enables proliferation at Palbociclib-exposed interfaces

Under Palbociclib treatment, human-like interfaces engaged a “mitochondrial tuning” programme encompassing histidine, folate, CoA and lipoate metabolism, enhancing mitochondrial function, redox resilience and energy production (**Fig. 6a**). These metabolites concentrated along tumour peripheries in treated brains, with cumulative abundance increasing by ∼15% compared to control (p = 0.017; **Fig. 6b**). In primary tumours, mitochondrial antioxidant genes *IDH2* and *MGST1* showed strong positive associations with proliferation (*MKI67*) and TP53 activity in spatial Xenium data, confirmed by DREMI-based dependency scores (**Fig. 6c**). Independent validation in a bulk medulloblastoma cohort revealed significant correlations between *IDH2* and *MKI67* (r = 0.55, p = 1.39 x 10^-27^) and *PCNA* (r = 0.602, p = 5.67 x 10^-34^), with similar trends for *MGST1* (**Fig. 6d**). Together, these findings indicate that interface tumour cells deploy mitochondrial and antioxidant programs to sustain proliferation despite partial CDK4/6 inhibition, mirroring the tight coupling between mitochondrial function and proliferative behaviour observed in human medulloblastoma.

**Fig. 6:**
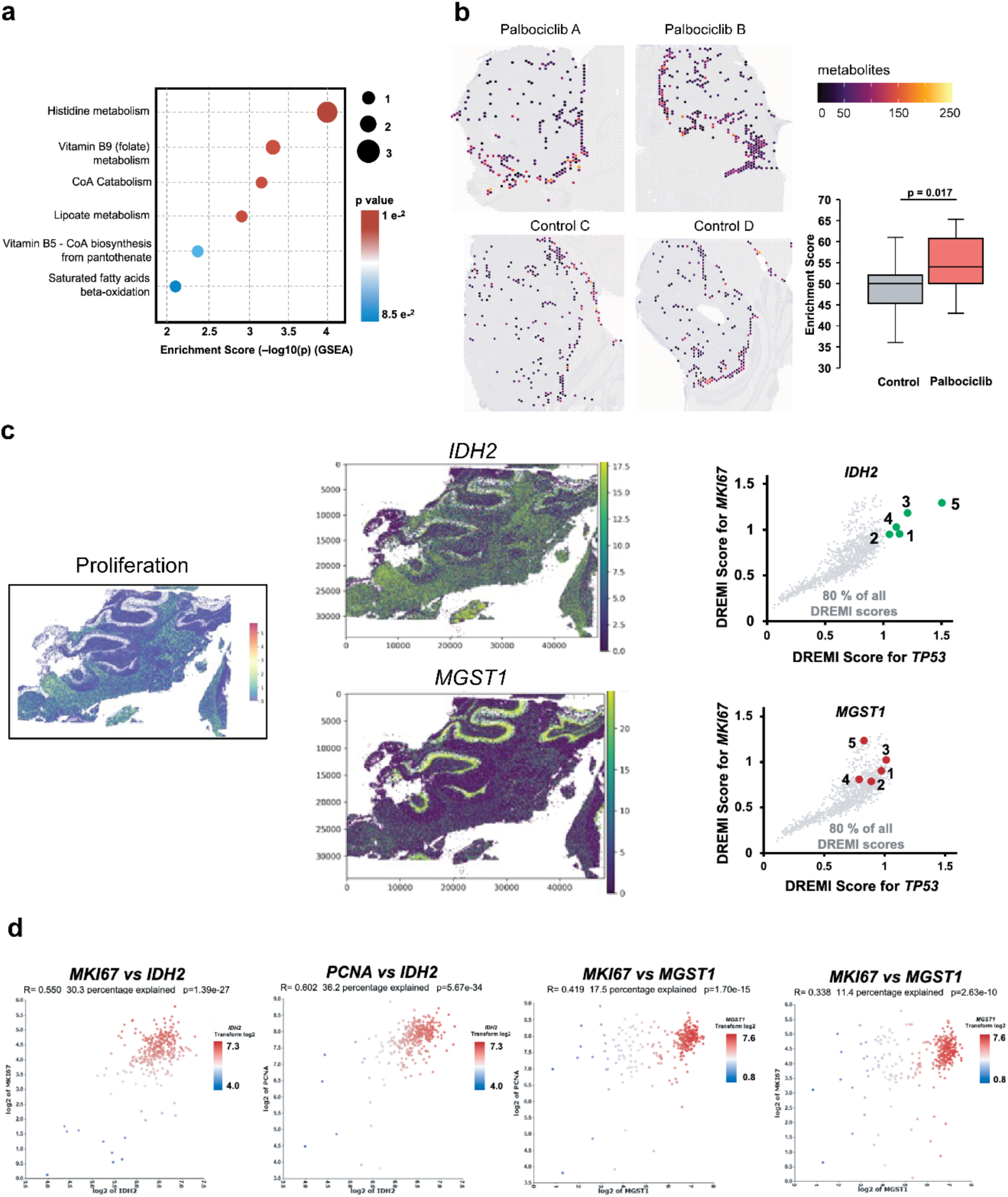
Mitochondrial tuning at the tumour–brain interface supports proliferation under Palbociclib treatment. **a** Metabolic pathways enriched in palbociclib-treated human-like interface cells. Pathway enrichment analysis of spatial metabolomics data comparing Palbociclib-treated versus control human-like interface regions in medulloblastoma PDOX brains. Dot plot shows pathways linked to mitochondrial function and redox homeostasis, including histidine metabolism, vitamin B9 (folate) metabolism, CoA catabolism and lipoate metabolism. Dot size indicates the number of altered metabolites and colour encodes the normalised enrichment score, highlighting a coordinated “mitochondrial tuning” signature in treated interface cells. **b** Spatial distribution of mitochondrial-tuning metabolites at the tumour - brain interface. Composite maps of metabolites belonging to mitochondrial-tuning pathways (histidine, folate, CoA and lipoate metabolism) are projected onto tissue sections from two Palbociclib-treated (Palbociclib A, Palbociclib B) and two control (Control C, Control D) PDOX brains. Human-like interface regions are outlined in red. Box plot quantifies the cumulative abundance of mitochondrial-tuning metabolites within the human-like interface, showing higher metabolite load in Palbociclib-treated versus control interfaces (15% increase; p = 0.017). **c** Spatial correlation of mitochondrial antioxidant genes with proliferation in human medulloblastoma. Xenium spatial transcriptomics from five human medulloblastomas show proliferative niches (left, MKI67 module score) and expression of mitochondrial antioxidant genes IDH2 and MGST1 (centre). Right, DREMI-based association analysis across tumours demonstrates strong positive relationships between IDH2 or MGST1 and proliferation (MKI67) as well as TP53 activity, indicating that mitochondrial tuning genes are preferentially expressed in proliferating cancer cell populations. **d** Independent validation of mitochondrial tuning–proliferation coupling in human data. Scatter plots from an external medulloblastoma dataset show significant positive correlations between IDH2 expression and the proliferation markers MKI67 and PCNA, and between MGST1 and MKI67 or PCNA (Pearson r and p values indicated). Together, these findings support a model in which mitochondrial tuning at the tumour–brain interface enhances energy production and redox protection, enabling sustained proliferation despite low Palbociclib exposure.

### Rare Palbociclib-positive proliferating spots express a mitotic and chemoresistance gene programme

Spatial integration identified rare Palbociclib-positive spots retaining high E2F activity despite drug exposure (**Fig. 7a, b**). Transcriptomic profiling revealed a distinct programme combining mitotic pathways such as chromosome segregation and spindle checkpoint signalling, with chemoresistance-associated genes and long non-coding RNAs (**Fig. 7c, d**). Ferroptosis-inhibitory genes were also upregulated, suggesting protection from ROS-driven cell death. Notably, resistance-associated genes, including *CDC5A*, linked to poor survival in independent medulloblastoma cohorts, were enriched alongside other cell-cycle regulators such as *CDK1*, *AURKA*, and long non-coding RNAs implicated in therapy resistance (**Fig. 7e, f**). Together, these findings define a drug-induced adaptive state in which a small subset of tumour cells maintains proliferation under CDK4/6 inhibition by reinforcing mitotic machinery and activating chemoresistance and ferroptosis-protective programs.

**Fig. 7:**
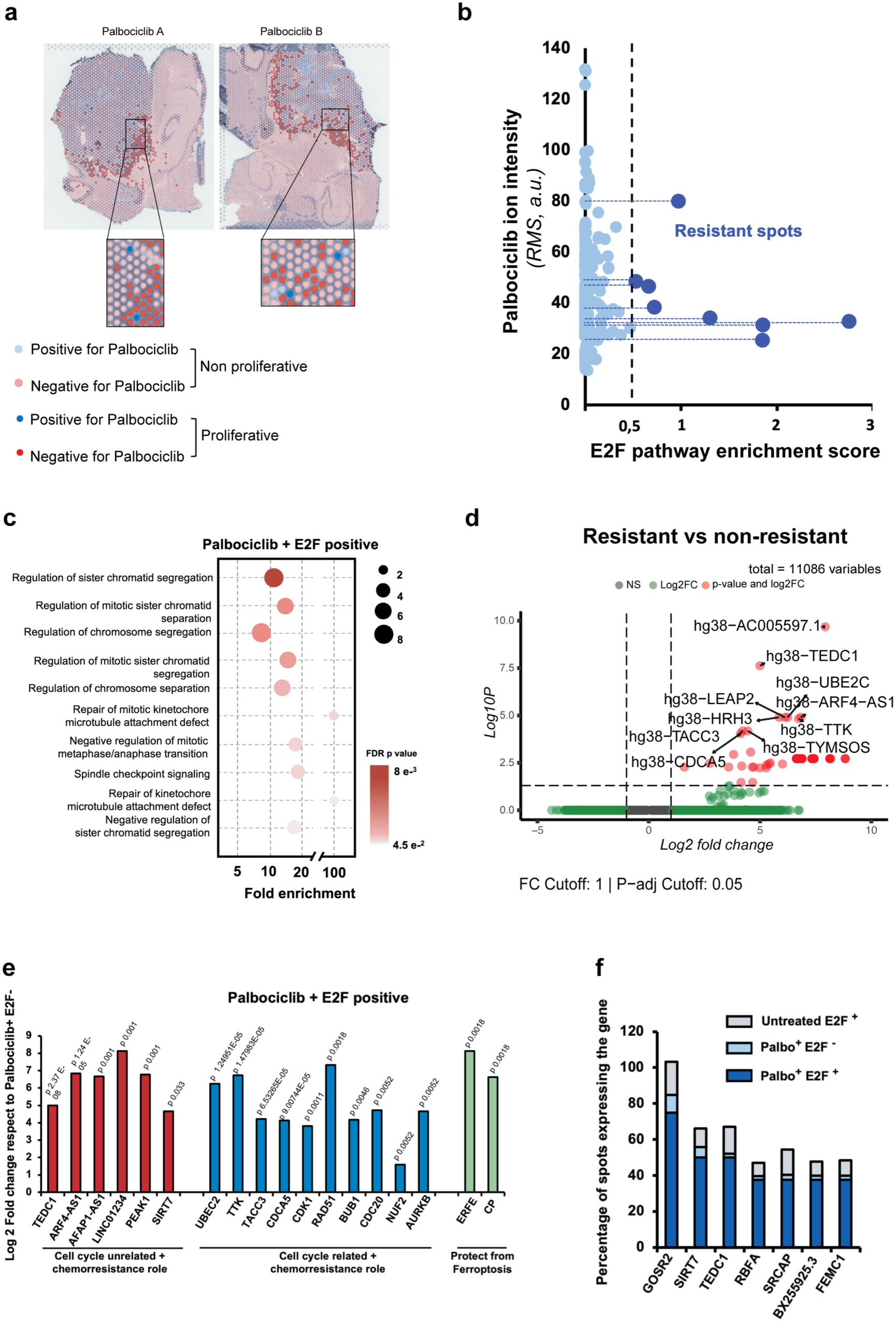
Spatially defined Palbociclib-resistant proliferative niches and associated gene programme. **a** Identification of Palbociclib-positive proliferating spots. MALDI–MSI–derived Palbociclib signal is overlaid on Visium spatial transcriptomic sections from two treated PDOX brains. Each Visium spot is classified according to drug exposure (Palbociclib-positive/negative) and and proliferative state (E2F-high/E2F-low). A small subset of Palbociclib-positive, E2F-high spots (blue) is detected within the tumour, indicating local proliferation despite drug exposure (“resistant niches”). **b** Relationship between Palbociclib signal and proliferative activity. Scatterplot of per-spot Palbociclib ion intensity (RMS, a.u.) versus E2F pathway enrichment score across all treated Visium spots. Most Palbociclib-positive spots show near-complete suppression of E2F activity across a range of Palbociclib levels, whereas a small set of eight Palbociclib-positive, E2F-high spots (“resistant spots”) maintain high E2F scores despite Palbociclib levels higher than those that abrogate proliferation elsewhere. **c** Pathways enriched in Palbociclib-positive proliferating niches. Gene ontology enrichment for transcripts upregulated in Palbociclib-positive, E2F-high spots (resistant) compared with Palbociclib-positive, E2F-low spots (non-resistant), highlighting enrichment of programmes related to mitosis and chromosome segregation. **d** Differential expression between resistant and non-resistant Palbociclib-positive spots. Volcano plot comparing Palbociclib-positive, E2F-high (resistant) versus Palbociclib-positive, E2F-low (non-resistant) spots. Numerous genes involved in cell cycle control and chromosomal segregation and chemoresistance-associated long non-coding and antisense RNAs are significantly upregulated in resistant spots. **e** Selected resistance-associated genes in Palbociclib-positive proliferating cells. Bar plots show log₂ fold changes for representative genes upregulated in Palbociclib-positive, E2F-high spots relative to Palbociclib-positive, E2F-low spots. Genes are grouped by functional category: cell cycle and chemoresistance (blue), chemoresistance-associated signalling and non-coding RNAs (red), and factors implicated in protection from ferroptosis (green), highlighting a multifaceted resistance programme. **f** Specific enrichment of resistance genes in Palbociclib-positive proliferating spots. For selected resistance-associated genes, stacked bars indicate the percentage of spots expressing each gene in untreated E2F-high, Palbociclib-positive E2F-low and Palbociclib-positive E2F-high (resistant) groups. Many genes are expressed in a large fraction of resistant spots but are rare in Palbociclib-positive non-proliferative and untreated proliferative regions, suggesting a Palbociclib-specific adaptive programme that sustains proliferation under therapeutic drug levels.

## Discussion

Single-cell and spatial transcriptomic studies have defined the developmental origins and intratumoural heterogeneity of SHH-MB and begun to reveal how targeted agents such as CDK4/6 inhibitors remodel tissue microenvironment^3,32,33^. Building on this foundation, we developed a same-section spatial pharmaco-multiomics framework that integrates spatial metabolomics with spatial transcriptomics to map palbociclib distribution, metabolites landscapes, and transcriptional states within the same SHH MB-PDOX and primary human tumour sections. In doing so, we confirm and extend previous findings that palbociclib suppresses E2F-driven proliferation and promotes neuronal differentiation across much of the tumour mass^3,8,9^, while demonstrating that residual drug tolerance is spatially concentrated at tumour–brain interfaces^3^.

Spatial multi-omics approaches are increasingly used to chart tumour ecosystems, yet most workflows either integrate RNA with protein or profile different modalities on adjacent sections, which can compromise spatial fidelity^13,34,35^. Although recent advances in mass spectrometry have enabled combined transcriptome and metabolite profiling on the same section, these efforts have largely focused on non-CNS tissues and typically lack direct drug measurements ^13,36^. Our pipeline combines same-section MALDI–MSI with Visium and high-resolution Xenium in matched regions, while preserving RNA quality and species-resolved annotation. By registering palbociclib intensity, metabolite distributions, and cell-state programs within the same histological context, we establish a generalisable spatial pharmaco-multiomics framework to dissect how local drug exposure, metabolism and transcriptional state intersect within microscale niches that remain invisible to bulk or dissociated profiling.

This framework reveals a sharp dichotomy between the drug-rich tumour core and the drug-poor tumour–brain interface. Consistent with Vo, et al. ^3^, palbociclib-rich tumour core undergoes a therapeutic shift towards neuronal differentiation: E2F targets and SHH-A proliferative signatures are suppressed, while neuronal differentiation and SHH-C programs are induced. By contrast, the tumour–brain interface and perivascular territories retain E2F-high proliferative states and are enriched for mesenchymal-like stromal populations and ECM-remodelling genes. Thus, proliferative reservoirs under CDK4/6 inhibition are anatomically constrained to microenvironments where palbociclib penetration is limited, likely shaped by ECM composition and BBB properties^9,37^. These findings provide a spatial explanation for high relapse rates after palbociclib withdrawal^8^ and underscore that optimising drug delivery to invasive margins is as critical as targeting intrinsic signalling pathways.

Spatial metabolomics adds a complementary layer by linking the proliferative territories to a ganglioside- and ECM-rich metabolic niche. Within human tumour regions, we identify a GM2- and sphingomyelin-enriched cluster associated with highly proliferative territories ^25–27^ and a phosphatidylcholine/phosphatidic-acid–dominated cluster associated with neuronal differentiation, synaptic programs, and higher palbociclib exposure^28,29^. GM2-high territories localise to tumour interfaces and invasive fronts where drug is scarce, and are enriched for ECM-associated metabolites, while differentiated, drug-rich bulk exhibits glycolipid patterns consistent with neuronal maturation^30,31^. These observations align with prior evidence linking GM2 and its synthesising enzyme B4GALNT1 to medulloblastoma aggressiveness and therapeutic vulnerability^38–40^. Moreover, the mesenchymal-like stromal populations and ECM–integrin interactions we detect at the interface echo “invasive edge ecosystems” in glioblastoma, where ECM remodelling, integrin signalling, and reactive gliosis drive invasion and immune exclusion^41–43^. Together, our data support a model in which glycosphingolipid–ECM cooperativity and mesenchymal–stromal crosstalk defines a permissive, palbociclib-poor niche for continued tumour expansion.

Beyond physical barrier effects, we uncover a distinct metabolic adaptation in therapy-exposed border cells. Modality-integrated pathway analysis reveals coordinated upregulation of histidine, folate, CoA and lipoate metabolism in the human-like interface under palbociclib treatment, a “mitochondrial tuning” program that supports oxidative phosphorylation, fatty-acid β-oxidation, redox buffering and anaplerosis. One-carbon and folate-linked pathways have been implicated in glioma growth, glioma stem cell survival and therapy resistance, and are being explored as metabolic vulnerabilities^44–46^. Here, we localise these pathways to a defined interface niche and link them to residual proliferative states under CDK4/6 inhibition. The spatial coupling between this metabolic signature, cardiolipin enrichment, and mitochondrial antioxidant genes such as *IDH2* and *MGST1*, validated in primary medulloblastomas and an independent bulk cohort, suggests that enhanced mitochondrial and redox capacity allows interface cells to sustain proliferation under submaximal drug exposure, consistent with adaptive CDK4/6 resistance in other solid tumours^47,48^. These data nominate mitochondrial tuning and one-carbon metabolism as testable combination targets at the tumour margin.

Finally, we identify rare palbociclib-positive, E2F-high “resistant spots” enriched for mitotic and chromosomal-segregation genes, highlighting an additional, orthogonal mode of resistance that cannot be attributed to drug scarcity alone. In breast cancers and some other cancers, CDK4/6 inhibitor resistance has been linked to alterations in the RB/CDK4/6–cyclin D axis, CDK2/cyclin E upregulation, PI3K–mTOR activation and mitochondrial rewiring ^49,50^. In SHH-MB, we instead observe spatially isolated populations that maintain high E2F activity at therapeutic palbociclib levels and upregulate mitotic checkpoint kinases, DNA repair factors, and chemoresistance-associated lncRNAs, together with ferroptosis-suppressive genes. This programme is largely absent from untreated proliferative regions and non-proliferative drug-exposed spots, suggesting a drug-induced adaptive state. These resistant cells may seed durable relapse and point to mitotic checkpoint, DNA repair and ferroptosis pathways as rational partners for CDK4/6 inhibition.

Interpretation of these findings is bounded by the models and measurements used. Our analyses were performed in an SHH MN-PDOX model in immunodeficient mice and a limited number of primary SHH tumours, which may not fully capture patient heterogeneity. MALDI-MSI provides relative metabolite coverage and is constrained by ionisation efficiency and isomeric complexity. Additionally, our approach infers spatial transitions from static profiles rather than tracking dynamic changes, and resolution mismatches between Visium, Xenium, and MALDI-MSI introduce within-spot heterogeneity. These factors mean that relationships between GM2-rich/ECM niches, mitochondrial tuning and resistant cells remain correlative, and their causal roles in long-term resistance are not yet proven.

Future work should extend this same-section framework to larger, molecularly diverse and longitudinal patient cohorts, including relapse material and additional brain tumour types. Incorporating complementary modalities such as proteomics, immune profiling and live-imaging readouts will provide deeper mechanistic insight. Targeted in vivo perturbations of ganglioside/ECM pathways, one-carbon and mitochondrial metabolism, and key resistance genes and lncRNA modules will be critical to test whether disrupting these spatially defined niches can improve CDK4/6-based treatment outcomes. These spatial observations may also inform current medulloblastoma local-control strategies that prioritise gross total resection and dose intensification to the tumour bed, by highlighting the tumour–brain interface as a reservoir of residual proliferative potential under therapy.

In conclusion, our same-section spatial pharmaco-multiomics approach reveals how SHH-MB responds to and resists CDK4/6 inhibition within spatially distinct niches. Palbociclib-rich tumour bulk undergoes neuronal differentiation, while drug-poor interfaces and rare palbociclib-positive resistant spots sustain proliferation through metabolic and mitotic adaptations. These findings argue for spatially informed combination strategies targeting both interface-specific metabolic liabilities and tumour-intrinsic resistance programs, providing a blueprint for precision therapies in medulloblastoma and other solid tumours.

## Methods

### MB-PDOX mouse model, Palbociclib treatment, and sample collection

The SHH MB-PDOX mouse model was established using seven- to nine-week-old male NSG mice. This model originates from a 4.9-year-old male patient with SHH-MB displaying desmoplastic/nodular morphology. Developed by the Olson laboratory at the Seattle Children’s Research Institute as part of the Children’s Oncology Group brain tumour biology study ACNS02B3, the model is publicly available through the Fred Hutchinson Cancer Research Center Biorepository (https://research.fredhutch.org/olson/en/btrl.html) and was previously characterised by Brabetz, et al. ^51^. All mice were housed under standard conditions, including a 12-hour light/dark cycle with ad libitum access to food and water, in compliance with NIH guidelines and the approvals of The University of Queensland Molecular Biosciences Animal Ethics Committee (IMB/386/18) and Institutional Human Research Ethics Committee (2015001410).

The establishment, treatment, and collection of PDOX mice were conducted using the same procedure described by Vo, et al. ^3^. For orthotopic xenograft implantation, mice were anesthetised, and a scalp incision was made to expose the calvarium. Using a handheld 0.7-mm microdrill, a burr hole was created above the right cerebellar hemisphere. Tumour cells suspended in serum-free DMEM at a concentration of 50,000 cells/μL were injected stereotactically to a depth of 2 mm below the dura. The surgical site was sealed with SurgiFoam (Johnson & Johnson MedTech, NJ, USA) to prevent cell leakage, and the scalp was closed with surgical staples. Post-implantation, animals were randomly assigned to treatment groups, receiving either 100 mg/kg Palbociclib hydrochloride (Pfizer, NY, USA) daily for seven days or no treatment (vehicle-treated). Palbociclib was dissolved in 50 mM sodium lactate, pH 4. An equivalent volume of 50 mM sodium lactate, pH 4 was administered orally daily to mice in vehicle groups (from now called controls). Treatment efficacy was assessed by monitoring drug response indicators and intratumoural heterogeneity. There are four PDOX mouse brains (2 Palbociclib-treated A,B and 2 controls C,D) for MALDI+Visium, and four SHH medulloblastoma PDOX mouse brains (2 Palbociclib-treated E,F and 2 controls G,H) for MALDI+Xenium.

### Human medulloblastoma FFPE samples

In total, five paediatric medulloblastoma samples were included in this study. Two SHH-MB samples, Human MB1 (ID: P59460) and Human MB2 (ID: P52407), were profiled using Visium, and their corresponding adjacent sections were analysed using Xenium. Three Group 3 medulloblastoma samples (IDs: P3868, P6055125 and P62560) were analysed using Xenium.

### Tissue sectioning

Tissue samples were sectioned using the Epredia™ CryoStar™ NX70 Cryostat (Thermo Fisher Scientific, MA, USA). Sections were cut at a thickness of 8 µm and mounted onto Visium glass slides or Xenium glass slides for MALDI-MSI processing, followed by Visium tissue optimisation or spatial gene expression analysis.

### Spatial metabolomics by MALDI-MSI

MALDI–MSI was performed directly on the same tissue sections used for Visium or Xenium, followed by H&E staining and library preparation (Visium) or *in situ* hybridisation (Xenium). Slides were vacuum-dried for 10 minutes and then scanned with a Braun FS-120 scanner and analysed using HistoView v1.00 software. Matrix application was performed using the SunCollect Sprayer (Sunchrom, Germany) at 2.5 psi nitrogen pressure, with a line distance of 2 mm, a z-position of 30 mm, and flow rates of 10 μL/min, 20 μL/min, and 40 μL/min for layers 1, 2, and ≥3, respectively. To detect metabolites, lipids, and drug distribution (Palbociclib) in positive ionisation mode, 12 layers of DHB (2,5-Dihydroxybenzoic acid) matrix (Sigma-Aldrich, MO, USA) were applied at a concentration of 30 mg/mL in ethanol.

Mass spectrometry imaging (MSI) was performed using a TIMS TOF Flex-MALDI-2 instrument (Bruker Daltonics, Germany) operated in M1 mode. Laser power was optimised at the start of each analysis and was typically set to 70–75%. Instrument calibration was carried out externally using the ESI-L tune mix (Agilent G1969-85000) in ESI mode. The instrument was operated with tims Control 4.1.13, Flex Imaging 7.2, Data Analysis 6.2, and Flex Imaging Batch Runner. All acquisition methods utilised a single-beam laser with a beam scan set to 46 μm (X and Y axes), resulting in a 50 μm pixel size, and 300 laser pulses per pixel at 10,000 Hz. The mass-to-charge ratio (m/z) range was 160–3000, with instrument parameters including isCID energy at 0 eV, ion energy at 5 eV, collision energy at 10 eV, and focus enabled. High-resolution captures were performed with a 20 μm pixel size. The MALDI-MSI acquisition parameters used were as follows: MALDI Plate Offset was set to 50 V, Deflection 1 Delta to 70 V, Funnel 1 RF and Funnel 2 RF to 350 Vpp each, and Multipole RD to 350 Vpp. The Collision RF ranged from 1000 to 4000 Vpp, while the Low mass (m/z) was set at 100. The Transfer Time was between 70 and 150 μs, and the Pre Pulse Storage was set at 10 μs.

MSI data were recalibrated post-acquisition using the appropriate matrix ion as a lock mass in Data Analysis software. The recalibrated dataset was normalised using the root mean squared (RMS) normalisation method in SCiLS Lab2024a (Bruker Daltonics). The spatial distribution of specific m/z values corresponding to analytes of interest was visualised and exported as TIFF files for further analysis.

MALDI–MSI was run at two resolutions (50 µm on Visium slide and 20µm on Xenium slide). Following the MSI scan, tissue sections on glass slides were washed twice in pre-chilled methanol for 30 seconds and stored at -80 °C. These samples were subsequently utilised for Visium tissue optimisation, gene expression, or Xenium spatial transcriptomics profiling experiments.

#### Spatial transcriptomics by Visium and Xenium

##### Tissue Optimisation

Tissue permeabilisation for Visium Spatial Transcriptomics was optimised according to the Visium Spatial Tissue Optimisation User Guide Rev E (10X Genomics, #CG000238), with specific modifications tailored to the unique characteristics of PDOX samples, which include both human tumour and mouse brain cells, as described by Vo, et al. ^3^. To ensure compatibility with spatial metabolomics data acquisition via MALDI-MSI, initial protocol development focused on optimising tissue processing on Visium slides. Permeabilisation timing was then carefully adjusted to enhance mRNA preservation following metabolomics analysis. These optimisations were validated through successful visualisation of the mouse cerebellar cortex structure and the identification of distinct metabolites in the white matter and molecular layers using MALDI-MSI **(Supplementary Fig. S1a)**. Subsequent H&E staining and Cy3-labeled mRNA signal analyses confirmed both the preservation of mRNA integrity and its uniform capture across the Visium slide, demonstrating the success of the optimised protocol **(Supplementary Fig. S1a)**.

##### Visium Spatial Gene Expression Library Preparation

Library preparation for sequencing was performed according to the 10X Visium Gene Expression User Guide (CG000239, Rev F) and the methodology described by Vo, et al. ^3^. PDX tissue sections, 8 μm in thickness, were mounted onto Visium slides based on conditions determined during the tissue optimisation phase. Permeabilisation was carried out for 8 minutes, followed by reverse transcription and cDNA amplification for 17 cycles. Indexed libraries were generated with 14 amplification cycles. The prepared libraries were pooled to a concentration of 1.8 pM and sequenced on a NextSeq2000 platform (Illumina). Details on sequencing data processing and read mapping were consistent with the protocols outlined in the Visium Gene Expression User Guide (CG000239, Rev F) and Vo, et al. ^3^.

##### Xenium spatial transcriptomics

Tissue sections stored at -80°C were processed for Xenium spatial transcriptomics following the Xenium *In Situ* for Fresh Frozen Tissues – Fixation & Permeabilisation User Guide (CG000581). Initially, the slides were incubated at 37°C for 1 minute, then immersed in a fixation solution for 30 minutes. A series of washes with 1X PBS (AM9624) and 1% SDS (71736) was performed, followed by permeabilisation in pre-chilled 70% methanol (34860) on ice for 60 minutes. Slides were then assembled into Xenium cassettes with PBS-T added for subsequent processing.

Probe hybridisation, ligation, and amplification were conducted according to the Xenium *In Situ* Gene Expression – Probe Hybridisation, Ligation & Amplification User Guide (CG000582). Tissue mRNA was targeted using the Xenium Human Brain Gene Expression Panel and hybridised at 50°C overnight. Unbound probes were removed, and a ligation step at 37°C for 2 hours joined the probe ends bound to target RNA, forming circular DNA molecules. These ligated probes were amplified to produce gene-specific barcodes. Fluorescently labeled oligonucleotides were then attached to the amplified DNA probes, and autofluorescence quenching was applied to minimise background noise. Tissue nuclei were stained with DAPI (4’,6-diamidino-2-phenylindole) to identify regions of interest. Finally, the tissue sections were loaded into a Xenium Analyzer for imaging and decoding, following the Xenium Analyzer User Guide (CG000584).

Despite the additional steps required for MALDI–MSI, our control experiments showed that the spatial transcriptomic data quality remained high (**Supplementary Figs. S1, S11**). When compared with previously generated Visium-only data from Vo, et al. ^3^ the multi-omics dataset showed nominally higher median UMIs (2,574 vs. 2,274) and gene counts (1,571 vs. 1,360), with only modest reductions in mean UMIs and gene counts (2,868 and 1,633 vs. 3,328 and 1,647), suggesting that MALDI–MSI and associated processing do not compromise spatial transcriptomic performance.

#### Post-Xenium hematoxylin –and eosin (H&E) staining

After Xenium analysis, tissues underwent H&E staining as per the Xenium *In Situ* Gene Expression – Post-Xenium Analyzer H&E Staining Guide (CG000613). Quencher removal was achieved with a Quencher Removal Solution, followed by Milli-Q water washes. Hematoxylin staining (MHS16) was performed, followed by bluing in CS702 and subsequent dehydration with ethanol (E7023). Eosin staining (3801615) was completed with additional ethanol and xylene (214736) washes. Finally, slides were cover-slipped using mounting media, dried for 30 minutes, and prepared for imaging.

### Mapping and validation of Palbociclib by MALDI–MSI

Palbociclib was prepared at a concentration of 1 μg/mL in 100% methanol. A 0.5 μL aliquot of this solution was mixed with an equal volume of matrix solution (30 mg/mL DHB in ethanol). The mixture was spotted onto a MALDI plate, where cocrystals were formed and analysed using MALDI-TOF in both MS1 and MS2 modes. To confirm the signal, wildtype male NOD.Cg-Prkdcscid Il2rgtm1Wjl/SzJ (NSG) mice were used for the experiment. The brains were collected and embedded, following the procedure described earlier for tumour collection. Tissue sections were then cryogenically sliced at 8 µm onto Superfrost microscope slides using a cryostat, as previously detailed. Palbociclib, in various concentrations (5 µg/mL, 2.5 µg/mL, 1.25 µg/mL, 0.25 µg/mL, and 0.025 µg/mL), was applied in 0.5 µL aliquots onto the cycled spot on a slide. Additionally, 0.2 µL of the drug solution at concentrations of 10 µg/mL, 5 µg/mL, and 2.5 µg/mL was separately dropped onto three different tissue sections. Subsequently, the tissue sections were spray-coated with DHB (30 mg/mL in ethanol) using the method previously mentioned. Finally, MALDI-MSI was performed according to the procedure described above.

MALDI–MSI specifically detected Palbociclib at m/z 447.2385 in treated, but not control brains. The tandem MS fragmentation confirmed this assignment by identifying characteristic fragment ions at m/z 450.2550 and 448.2399 (**Supplementary Fig. S4**). Validation experiments on spiked glass slides and NSG mouse tissues established the specificity and dynamic range of Palbociclib detection (**Supplementary Fig. S4**). In MB-PDOX sections, Palbociclib signal at these m/z values was exclusively observed in treated samples (**Fig 2; Supplementary Fig. S4**).

### Spatial transcriptomics data analysis

Visium data processing and analysis: The overall Visium analysis process was conducted as described by Vo, et al. ^3^. This included read mapping, spatial stratification of spots by species and tissue regions, bioinformatics analysis of gene expression markers, differential expression analysis gene set enrichment analysis (GSEA), and the calculation of gene set enrichment scores for individual spots using transcriptional signatures generated by scRNA-seq of SHH MB patient samples for both PDOX model and human samples, and using transcriptional signatures generated by GSEA of PDOX for human samples. Clustering and cell-cell interactions were analysed by stLearn^52^. Besides unsupervised mapping of spots, expert annotation of H&E images from Visium and Xenium tissues was provided by Dr Thomas.

Xenium data processing and analysis: Quality control was performed on Xenium datasets for both human and mouse PDOX samples. Low-quality cells were filtered based on a minimum threshold of three detected genes per cell. Data integration was performed separately for human and mouse datasets using the Harmony^53^ algorithm to correct for batch effects. Following integration, unsupervised clustering was performed on each dataset independently using the Louvain algorithm implemented in Scanpy^54^. Cell type annotation was subsequently performed using key marker genes annotation from the 10x Genomics Human Brain Gene Panel. To investigate spatial cellular communication, cell-cell interaction analysis was performed using the grid mode implementation within the stLearn^52^ software.

### Multimodal integration of MALDI-MSI and Spatial Transcriptomics

Registration of metabolite signals from MALDI_MSI to spatial transcriptomics platforms was performed to enable multimodal analysis. For Visium data, alignment followed the SpaMTP manual registration workflow, mapping 55 µm diameter Visium spots (100 µm center-to-center distance) to the 50 µm continuous pixel resolution of the MALDI-MSI data. For Xenium data, metabolite signals (20 µm pixel resolution) were aligned using image-based affine transformation registration via the SimpleITK^55^ to enable the detailed alignment. The final aligned datasets were stored within the SpaMTP objects for visualization and converted to stLearn objects for downstream cell proportion analysis.

### Metabolomics analysis

Post-acquisition, mass spectrometry imaging (MSI) data were recalibrated using the appropriate matrix ion as a lock mass in Bruker’s Data Analysis Software (v6.1). The recalibrated dataset was imported into SCiLS Lab MVS 2024a, where it was normalised using root mean squared (RMS) normalisation. Spatial distributions of specific m/z values corresponding to analytes of interest were visualized and exported as TIFF files. Additionally, the raw MSI data were exported as .imzML files using SCiLS Lab for further downstream analyses using the developed SpaMTP^56^ pipeline. Additionally, the raw MSI data were exported as .imzML files using SCiLS Lab for further downstream analyses using the developed SpaMTP^56^ pipeline.

The metabolite annotation was performed using SpaMTP^56^ against database HMDB 5.0. Differential metabolite expression analysis across multiple samples was conducted using a pseudobulk aggregation strategy followed by edgeR. For comparisons between clusters, we utilised the FindAllMarkers function in Seurat with the Wilcoxon Rank Sum test. Unsupervised clustering of the metabolite profiles followed the standard Seurat workflow, utilising the top principal components (PCs) derived from the most variable metabolites. Finally, SpaMTP was employed to conduct pathway enrichment analysis and to visualise differentially expressed metabolites via heatmaps, volcano plots, and spatial tissue maps.

### Integrated spatial multiomics pathway analysis

To dissect the interface between invading tumour and healthy brain, we took advantage of two distinct spatial techniques. First, spatial transcriptomics was used to set two different compartments in the unique shared interface. Based on human gene enrichment, we set a boundary at 50 % enrichment in human transcripts. The interface composed by 50 to 90 % of human transcript enrichment was considered human-like interface (enriched in tumour cells), while the interface enriched in 50 to 10 % of human transcripts was set as mouse-like interface (enriched in mouse cells). Next, metabolites obtained using spatial metabolomics through MALDI were matched with human-like and mouse-like interface. Afterwards, DE metabolites in each compartment of the interface were obtained and pathway analysis was carried out. When comparing Control human-like and mouse-like, spaMTP was used. Comparison between Palbociclib-treated human-like and control human-like interface was carried out using Functional analysis function from MetaboAnalyst database (Positive Ion mode, T Score, Pathway library: Homo sapiens (human) [MFN], Mummichog and GSEA algorithm). The whole metabolite list was uploaded, as requested by the platform. Significantly upregulated metabolites were plotted in the Palbociclib-treated human-like and control human-like interface for further confirmation and visualisation. P value from Palbociclib-treated human-like and control human-like was calculated using 6 pseudoreplicates per sample (12 pseudoreplicates per group) using Wilcoxon rank-sum test.

Utilising the co-profiling of both metabolites and genes in the same tissue section, we applied multiple joint analysis approaches to produce new insights that unimodal data cannot. Joint pathway analysis combined both DE genes and DE metabolites into a network using the union of pathway maps from HMDB, reactome, kegg and wikiPathways, which comprehensively reports the links between genes and metabolites (e.g., binding, activation). To generate a joint ranking, relative abundances of genes and metabolites were normalised to a common scale using maximum value scaling. Pathway enrichment was subsequently assessed via GSEA^57,58^. The enrichment for the joint differentially expressed genes and metabolites increased the detection power of relevant pathways with supporting evidence from both modalities. For significantly enriched pathways associated with drug treatment, we visualised spatial distribution of maker genes and metabolites across the tissue section using three-dimensional plotting from SpaMTP, allowing for intuitive interpretation of the data.

### Integrated metabolomics and transcriptomics to map and characterise Palbociclib resistant cells

Resistant cells were characterised as Palbociclib positive and positive for E2F pathway genes (the E2F pathway score was calculated using AddModuleScore function for Seurat and the E2F pathway gene set from MSigDB), with a minimum enrichment score < 0.5 for the E2F pathway. This Palbociclib concentration threshold eliminates false positive signal in control samples, hence spots passed filtering are with high confidence. The same accounted for E2F pathway enrichment, removing any spots with E2F enrichment below 0.5, DEG between Palbociclib positive and E2F negative spots and Palbociclib positive and E2F positive spots were obtained using Seurat. GO enrichment analysis was carried out using DEG (GeneOntology Resource, p value cutoff 0.05) DEG between Palbociclib positive and Untreated Control spots were obtained using Seurat. All the genes found in at least one resistant spot were considered as DEG. DEG were further classified according to their function (Cell cycle in blue, chemoresistance in red, ferroptosis in green). based on existing references. The percentage of spots expressing the identified DEG between Palbociclib resistant spots and Palbociclib positive E2F negative spots were analysed in all the untreated control tumour spots. Only genes present in at least 37 % of resistant spots were analysed.

### Identifying related pathways in mouse PDOX models with biological process found in human MB

Further analysis was carried out to correlate the PDOX findings with human medulloblastoma data. Using scprep^59^, We obtained the knn-DREMI score (hereinafter DREMI score) for IDH2 and MGST1, two genes related with mitochondrial proper function, with the proliferation marker MKI67 and cancer marker TP53 in five human medulloblastoma samples using Xenium. Using scprep^59^, We obtained the knn-DREMI score (hereinafter DREMI score) for IDH2 and MGST1, two genes related with mitochondrial proper function, with the proliferation marker MKI67 and cancer marker TP53 in five human medulloblastoma samples using Xenium. The DREMI scores for all the 220 genes analysed by Xenium were calculated and considered “random” DREMI scores. Further validation was performed in a bigger patient cohort using publicly available data from R2: Genomics Analysis and Visualization Platform (http://r2.amc.nl http://r2platform.com). (Tumour Medulloblastoma - Williamson - 331 - tpm – ensh38e78: Source: ArrayExpress ID: emtab10767 Date: 2000-01-01), where we correlated IDH2 and MGST1 genes with proliferation markers MKI67 and PCNA.

## Supplementary figures

**Supplementary Fig. S1.**
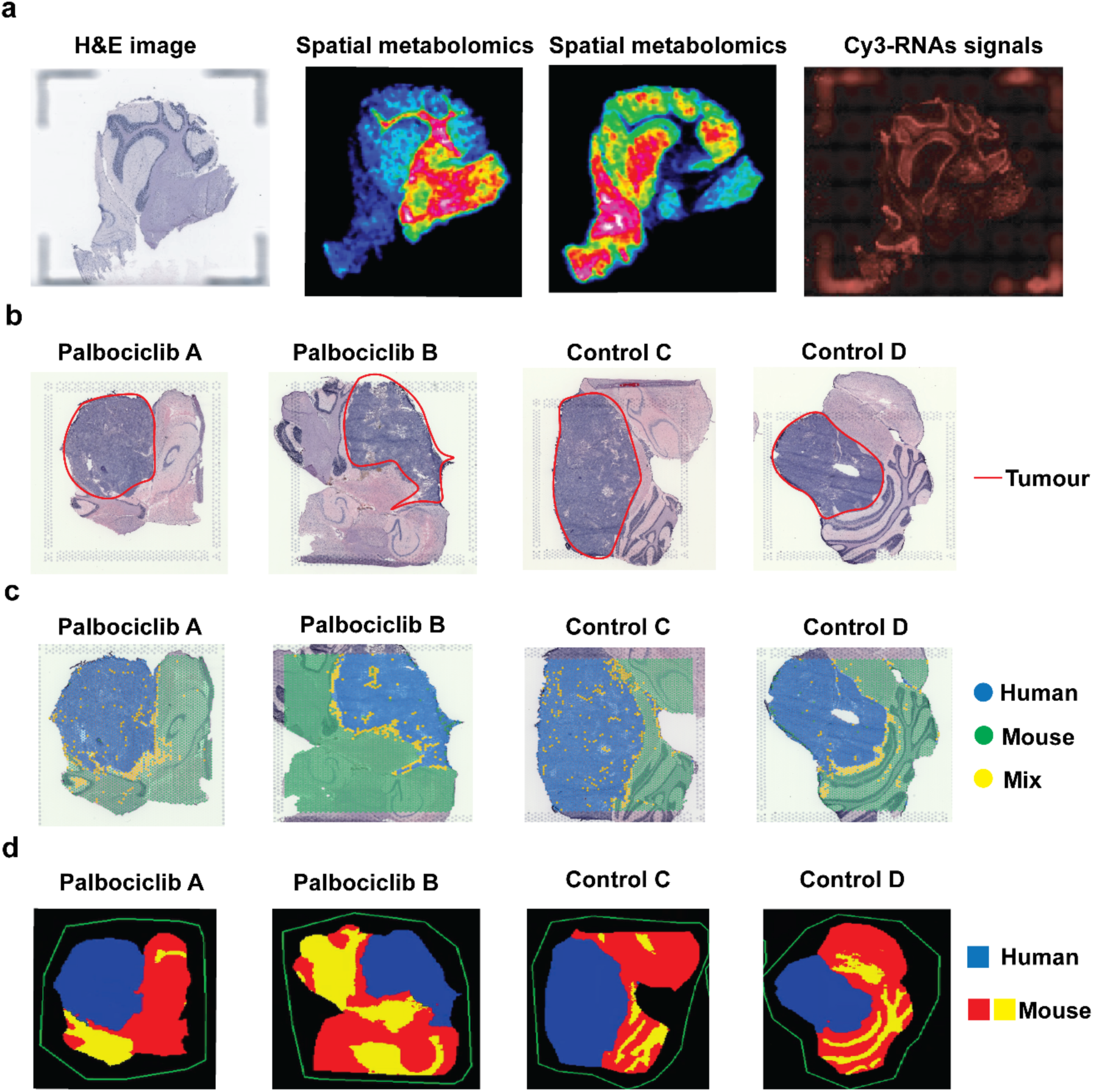
Optimisation and spatial annotation of human and mouse compartments in MB-PDOX brains. **a** Tissue optimisation. Representative H&E image, MALDI–MSI spatial metabolomics maps at two m/z values, and Cy3-RNA signal from Visium optimisation (8 min permeabilization), illustrating preserved morphology, metabolite distribution and robust spatial RNA capture in MB-PDOX sections. **b** Tumour delineation by H&E. H&E-stained sections from Palbociclib-treated (Palbociclib A, Palbociclib B) and control (Control C, Control D) MB-PDOX brains, with tumour regions outlined in red. **c** Species assignment of Visium spots. Visium spatial transcriptomic maps showing classification of spots as human tumour, mouse brain or mixed based on species-specific gene expression (blue, yellow and green, respectively). **d** Segmentation of spatial metabolomics data. Coarse-grained segmentation of MALDI–MSI data (50 µm) into human (blue) and mouse (red and yellow) regions following integration with Visium species labels, providing matched metabolic and transcriptomic annotation of tumour and host compartments.

**Supplementary Fig. S2.**
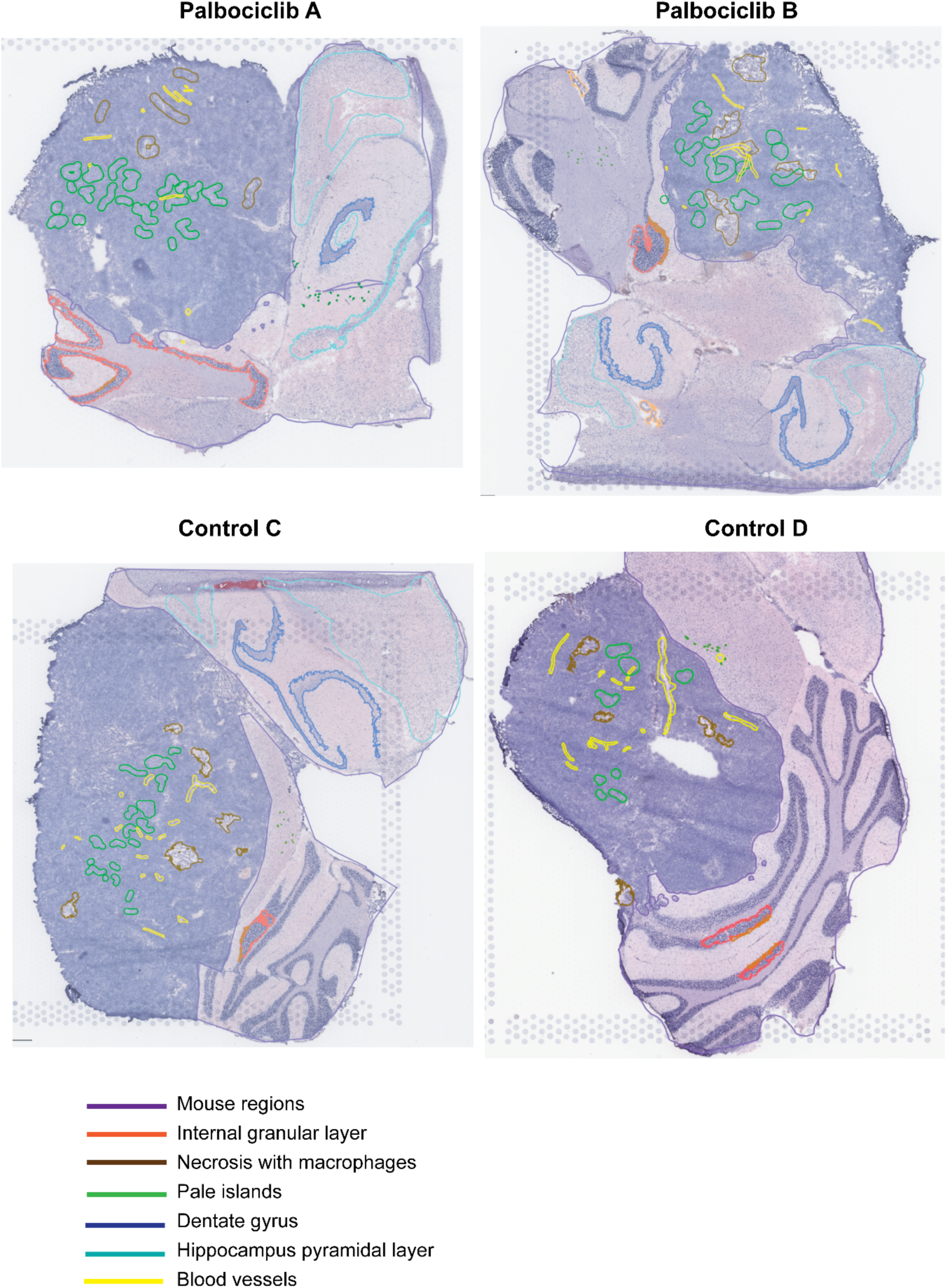
Histopathological annotation of Visium MB-PDOX samples. Representative H&E-stained sections from Palbociclib-treated (Palbociclib A, Palbociclib B) and control (Control C, Control D) MB-PDOX brains with manual annotations by a neuropathologist. Overlays delineate mouse brain regions, internal granular layer, areas of necrosis with macrophages, pale islands, dentate gyrus, hippocampal pyramidal layer and blood vessels, providing structural context for the spatial transcriptomic and metabolomic analyses.

**Supplementary Fig. S3.**
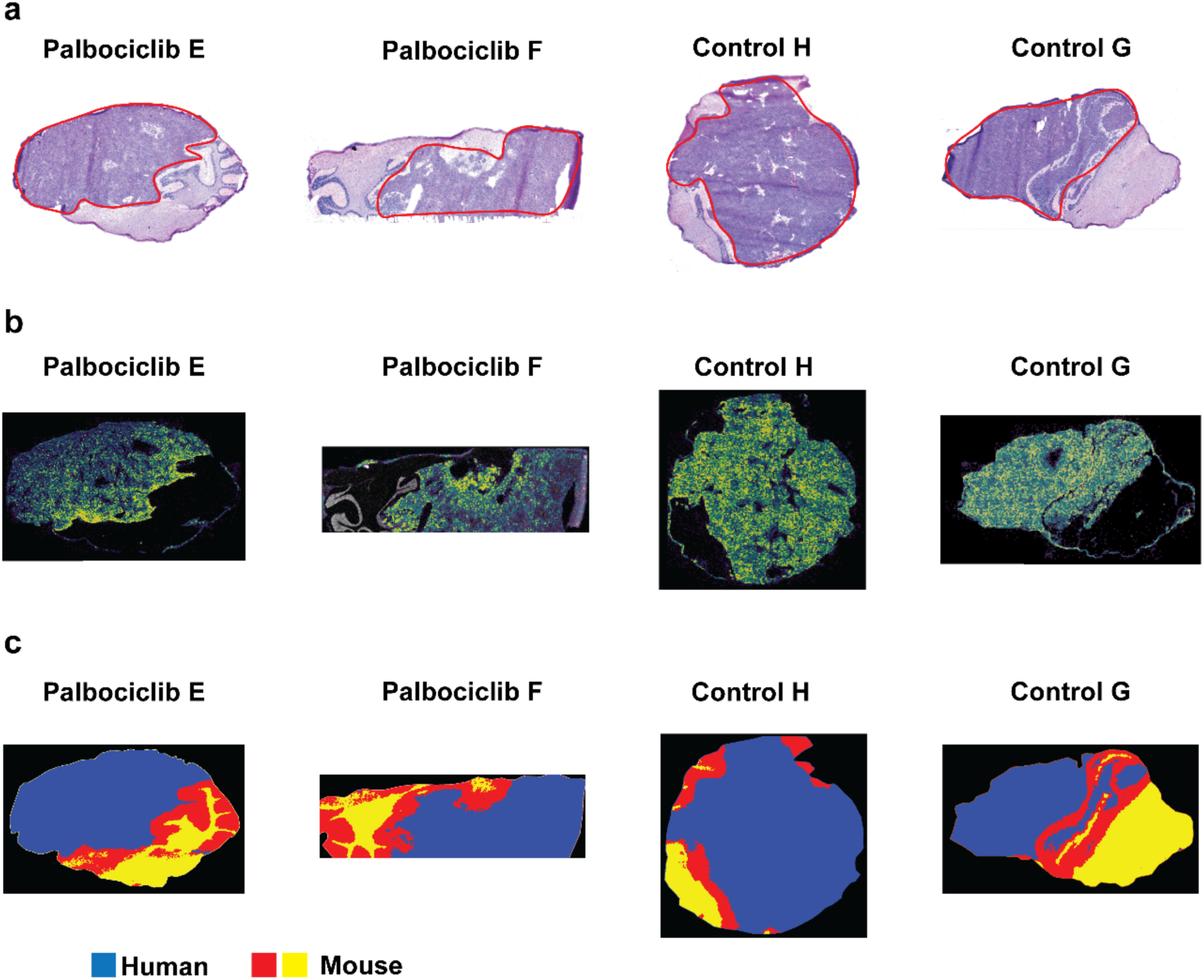
Single-cell–resolution annotation of human tumour and mouse brain in Xenium MB-PDOX samples. **a** H&E-stained sections from Palbociclib-treated (Palbociclib E, Palbociclib F) and control (Control H, Control G) MB-PDOX brains, with tumour regions outlined in red. **b** Xenium spatial transcriptomics from the same sections, showing single-cell transcript profiles within the outlined tumour areas. Signal is concentrated within human tumour regions and largely absent from surrounding mouse brain, enabling species- and tumour-specific mapping at cellular resolution. **c** Corresponding segmentation maps derived from Xenium species calls and co-registered MALDI–MSI at 20 µm, displaying human tumour (red) and mouse brain (green and yellow) compartments. These maps provide a refined view of tumour–host organisation for downstream single-cell and metabolomic analyses.

**Supplementary Fig. S4.**
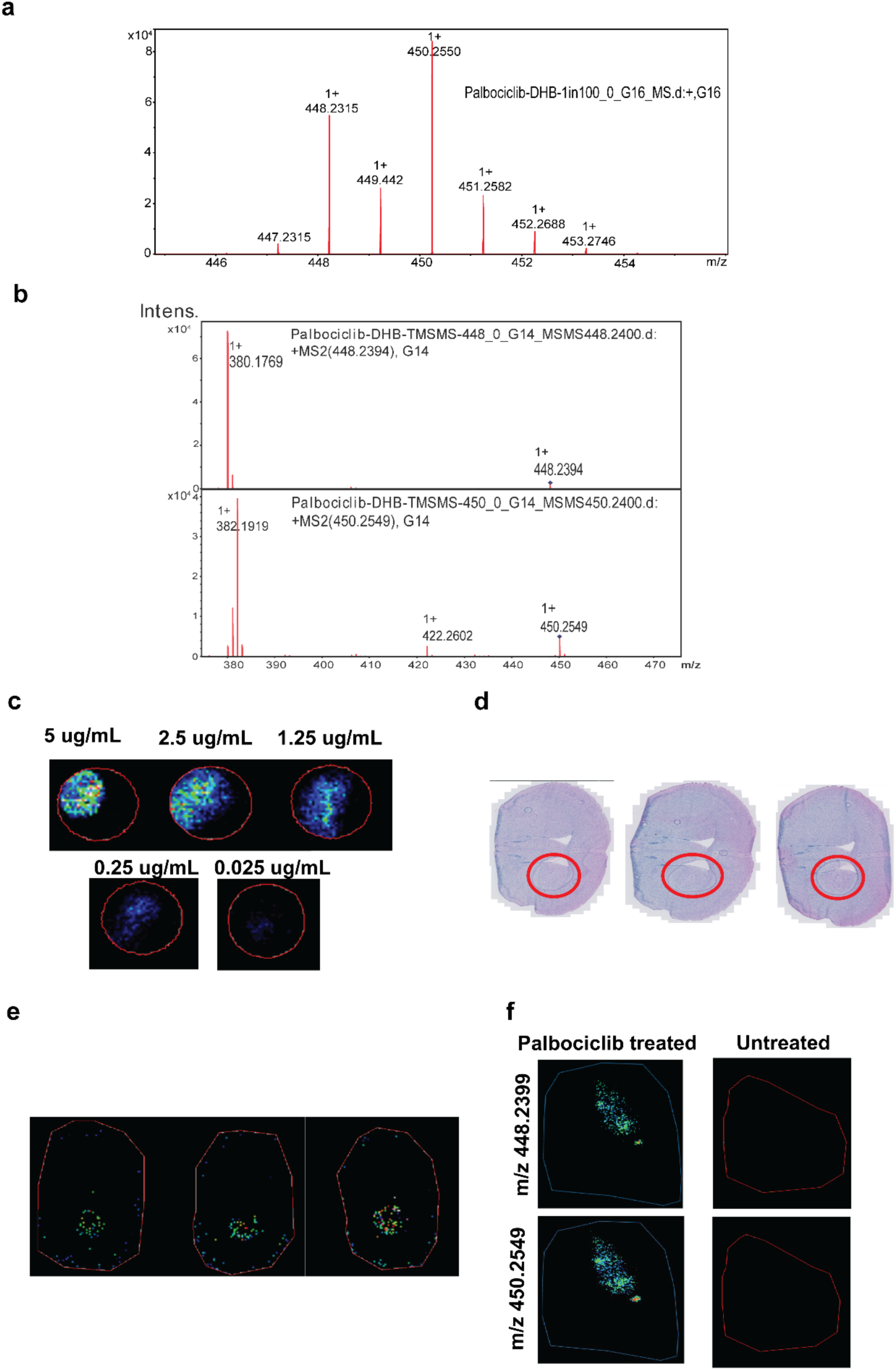
Validation of Palbociclib detection by MALDI–MSI. **a** MS¹ spectrum acquired by MALDI–MSI from Palbociclib-treated MB-PDOX tissue, showing the protonated Palbociclib ion and related adducts; the principal peak corresponds to the expected m/z of Palbociclib. **b** Reference MS² spectra obtained from a Palbociclib standard mixed 1:1 with 30 mg/ml DHB matrix in ethanol. The selected precursor ion (blue dot) and its characteristic fragment ions confirm the identity of the Palbociclib peak detected in tissue. **c** MALDI–MSI ion images of Palbociclib spotted on glass slides at decreasing levels (5, 2.5, 1.25, 0.25 and 0.025 µg/mL), demonstrating a robust dynamic range of drug detection. **d** H&E-stained sections from three wild-type independent mouse brains, with application sites indicated (red circles). **e** Corresponding MALDI–MSI maps of the mouse brains in e showing Palbociclib signal confined to the application area within each brain section. **f** Detection of Palbociclib in MB-PDOX brains after 3 days of systemic treatment versus untreated controls. MALDI–MSI ion images at m/z 448.2399 and 450.2549 confirm the presence of Palbociclib only in treated tumours and absence of signal in untreated tissue.

**Supplementary Fig. 5.**
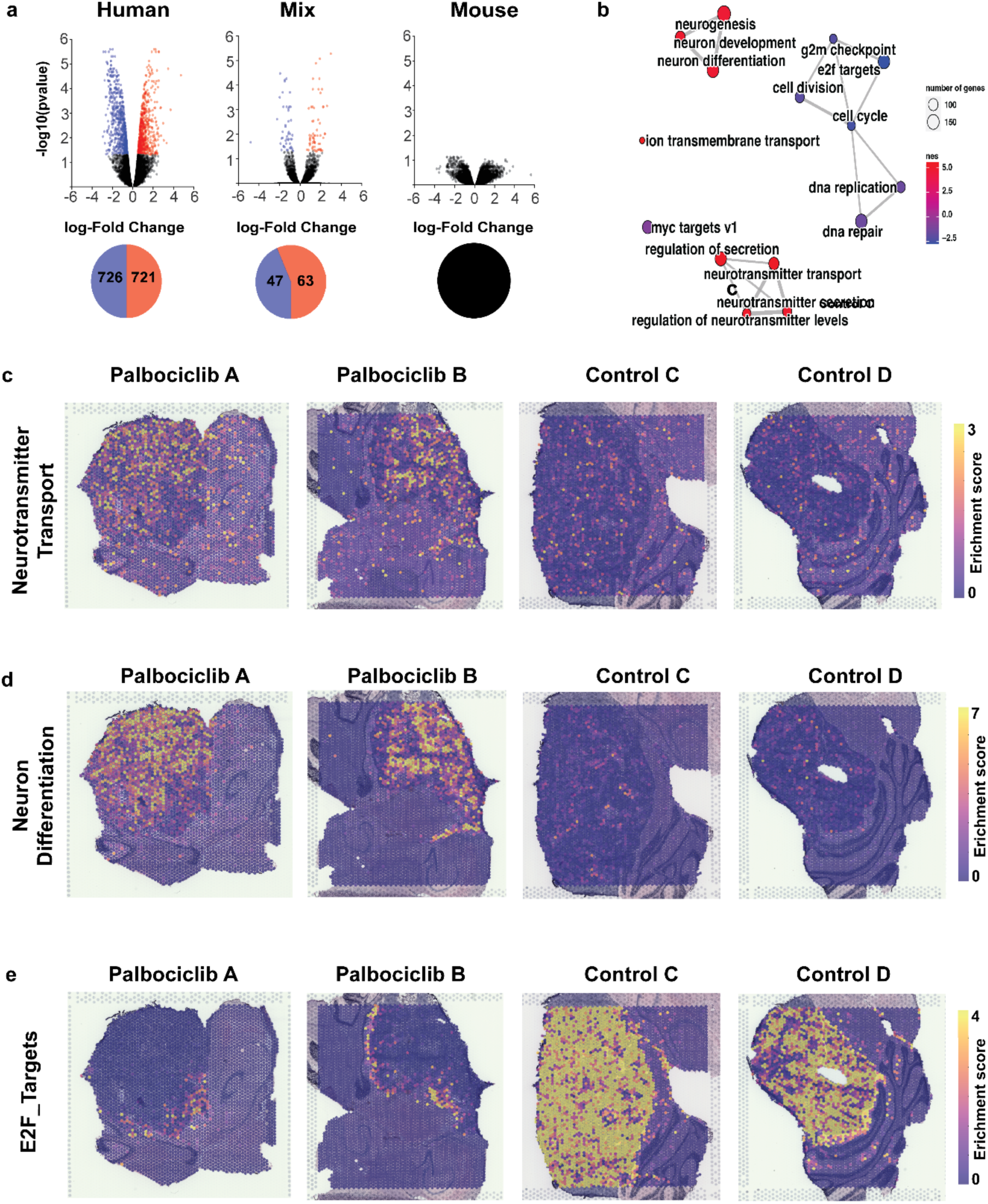
Spatial localisation of differentiation and proliferative programs in SHH MB-PDOX after Palbociclib treatment. **a** Differential expression by species compartment. Volcano plots of Visium spatial transcriptomics comparing Palbociclib-treated versus control tumours within human, mixed and mouse spot groups. In human spots, 726 genes are significantly upregulated and 721 downregulated; in mixed spots, 47 genes are upregulated and 63 downregulated. No genes reach significance in mouse spots, indicating that transcriptional remodeling is confined to the human tumour and mixed compartments. **b** Pathway enrichment in human tumour regions. Gene set enrichment analysis of differentially expressed genes in human spots identifies upregulation of neuronal differentiation and neurotransmitter transport pathways, and downregulation of E2F target and cell-cycle programs in Palbociclib-treated tumours. Human spots revealed upregulation of pathways associated with neuronal differentiation and neurotransmitter transport, consistent with a shift towards a less proliferative, more differentiated tumour cell state. In contrast, cell-cycle pathways, including E2F targets, were significantly downregulated, in line with Palbociclib’s role in inhibiting CDK4/6 activity and suppressing G1/S cell-cycle progression Dot size reflects the number of genes per pathway and colour denotes normalised enrichment score. **c–e**, Spatial maps of key gene programs. Visium enrichment scores for (c) neurotransmitter transport, (d) neuron differentiation and (e) *E2F* target gene sets across Palbociclib-treated (Palbociclib A, Palbociclib B) and control (Control C, Control D) sections. Warmer colours indicate higher pathway activity. Treated tumours show expanded regions with high neuronal differentiation and neurotransmitter transport scores and reduced *E2F* target activity, consistent with a shift from proliferative to differentiated states.

**Supplementary Fig. S6.**
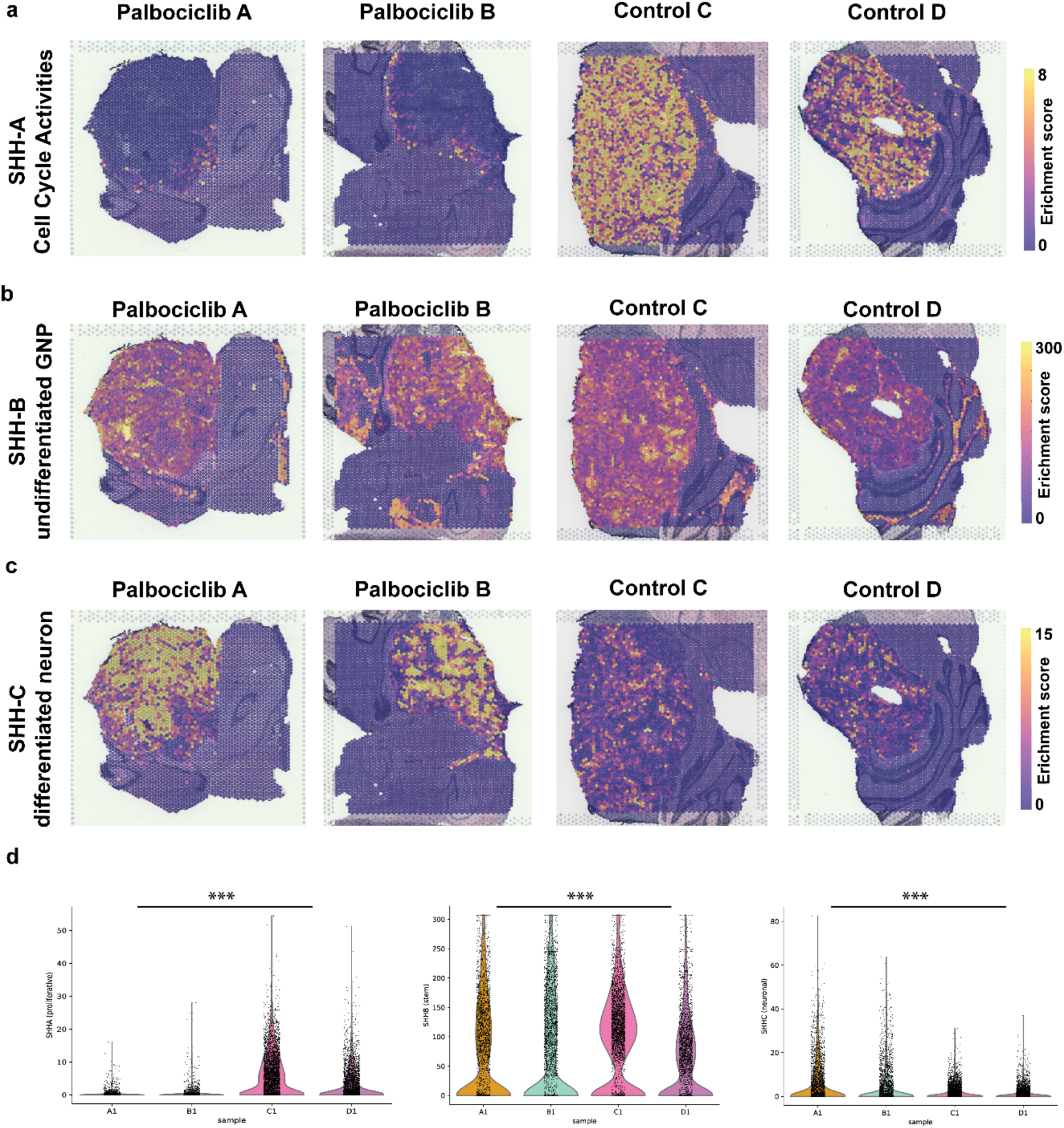
Spatial localisation of SHH medulloblastoma cell-state signatures after Palbociclib treatment. **a** Spatial maps of SHH-A (cell cycle–active) gene-set enrichment scores in Palbociclib-treated (Palbociclib A, Palbociclib B) and control (Control C, Control D) PDOX tumours. Spots are coloured by enrichment score, from low (purple) to high (yellow), illustrating a marked reduction of SHH-A activity in treated tumours. SHH-A signatures were diminished and spatially confined to the tumour interface in treated brains. **b** As in a, but for the SHH-B granule neuron progenitor (GNP-like) signature, showing redistribution of progenitor-like states with treatment. **c** As in a, but for the SHH-C differentiated neuron signature, revealing expansion of differentiated territories in Palbociclib-treated tumours relative to controls. SHH-C signatures were increased and formed discrete clusters within the tumour bulk. **d** Violin plots of per-spot SHH-A, SHH-B and SHH-C enrichment scores, grouped by sample. Asterisks indicate significant differences between treated and control tumours (P < 0.001, pseudosampling-based two-tailed t-test), confirming a global shift from proliferative SHH-A towards more differentiated SHH-C states following Palbociclib treatment.

**Supplementary Fig. S7.**
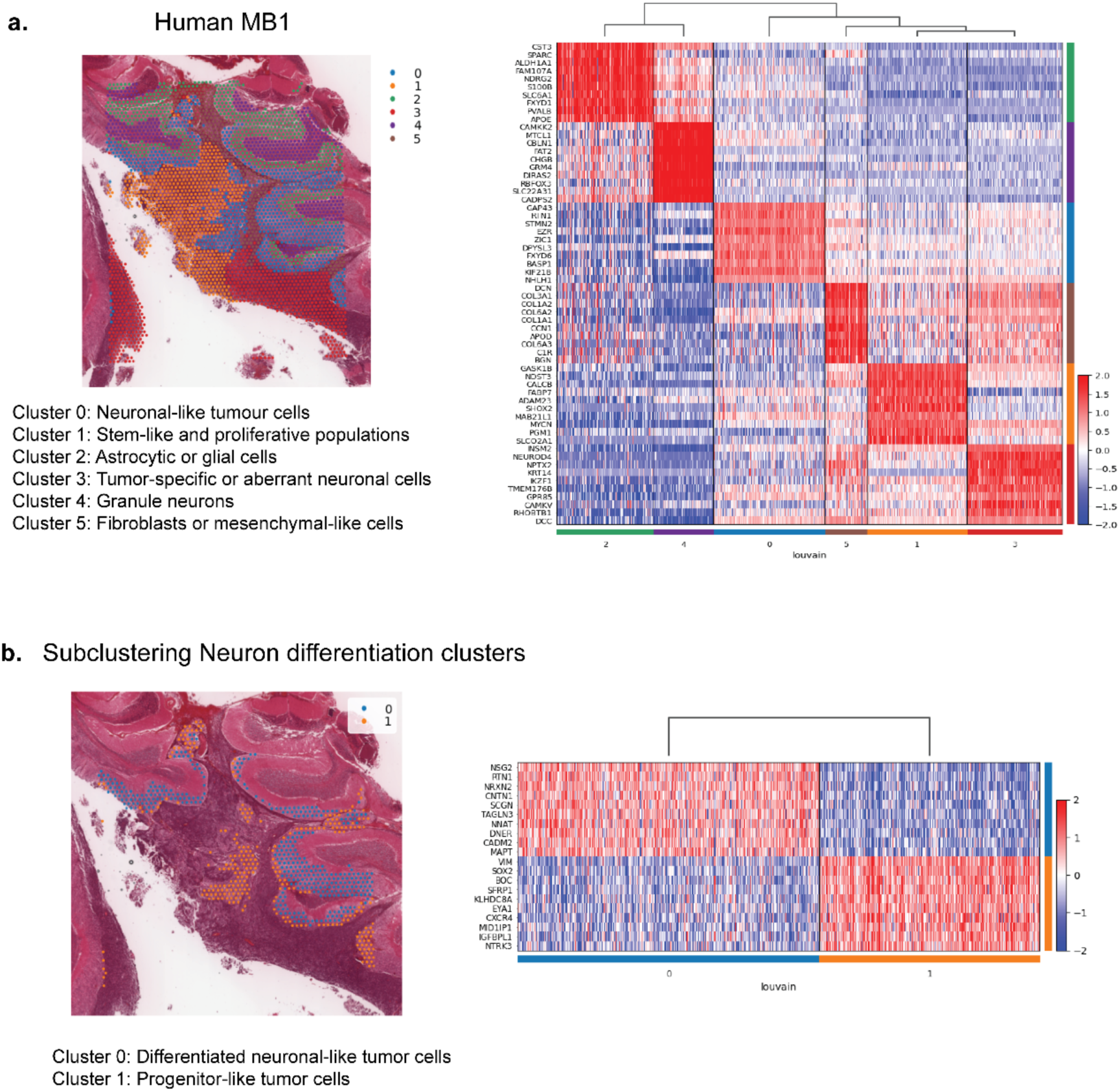
Spatial clustering of cell states in a human SHH medulloblastoma (Human MB1). **a** Global clustering of Human MB1. Unsupervised clustering of spatial transcriptomic spots from Human MB1 identifies six transcriptionally distinct populations. Left, spatial map showing the distribution of clusters across the tissue section. Right, heatmap of representative marker genes illustrating cluster-specific expression profiles. Cluster 0 corresponds to neuron-like tumour cells, Cluster 1 to stem-like and highly proliferative populations, Cluster 2 to astrocyte- or Bergmann glia–like cells, Cluster 3 to tumour-specific or aberrantly differentiated neuronal cells, Cluster 4 to cerebellar granule neurons, and Cluster 5 to fibroblast- or mesenchymal-like cells enriched for extracellular matrix genes, highlighting pronounced intratumoural and microenvironmental heterogeneity. **b** Subclustering of neuron-differentiation–related tumour cells. Further subclustering of the neuron-like tumour clusters (Clusters 0 and 3) resolves two major states. Left, spatial distribution of subclusters across the section. Right, heatmap showing that Subcluster 0 is enriched for neuronal differentiation markers, consistent with more differentiated neuron-like tumour cells, whereas Subcluster 1 expresses progenitor/stem-like and plasticity-associated genes, consistent with less differentiated, highly proliferative tumour populations.

**Supplementary Fig. S8.**
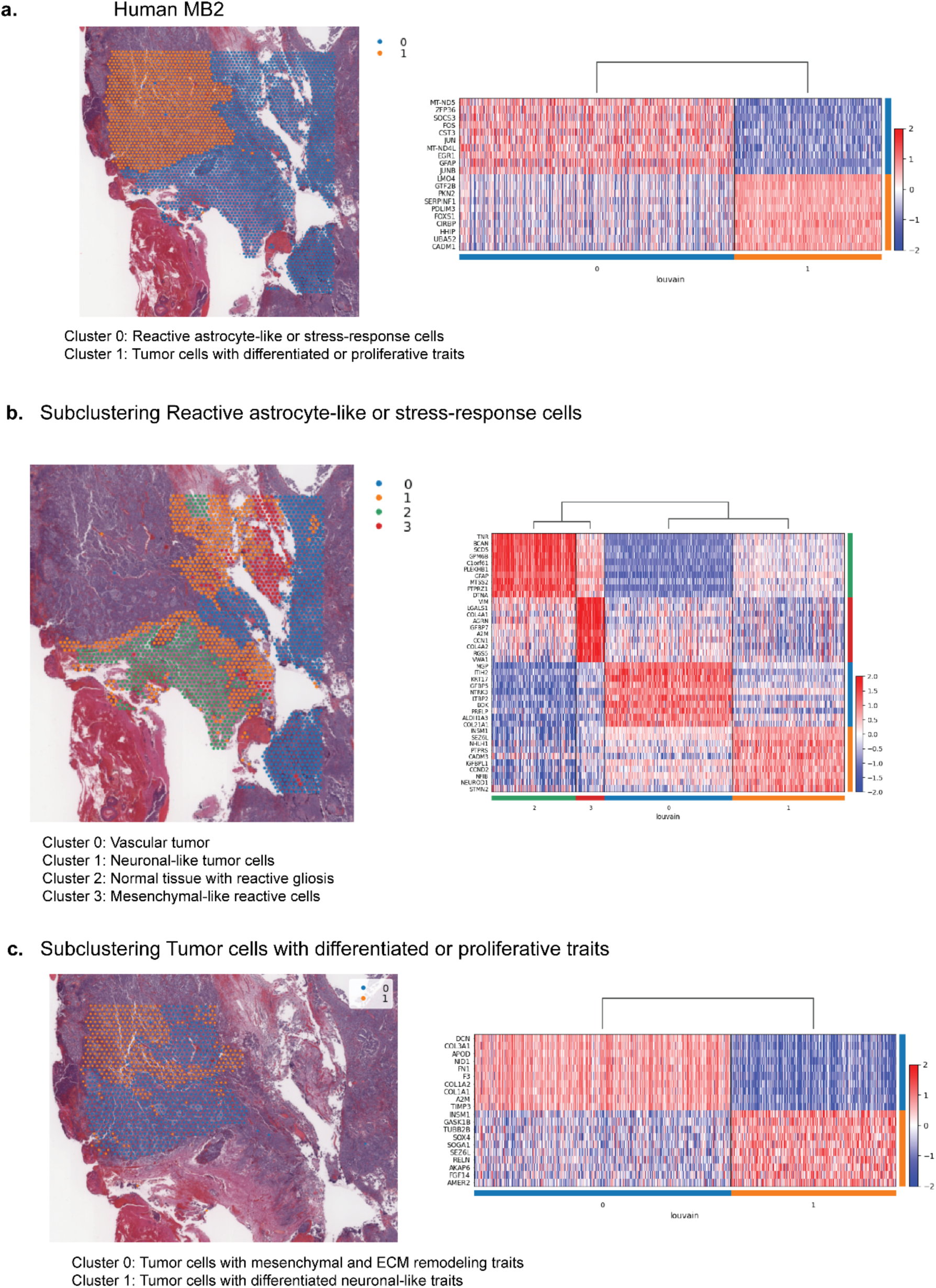
Spatial clustering of cell states in a second human SHH medulloblastoma (Human MB2). **a** Global clustering of Human MB2. Unsupervised clustering of spatial transcriptomic spots from Human MB2 resolves two major populations. Left, spatial map showing cluster distribution across the section. Right, heatmap of representative marker genes. Cluster 0 is enriched for astrocytic and stress-response markers (for example CST3, GFAP, FOS, JUN, JUNB), consistent with reactive astrocyte-like or stress-response cells. Cluster 1 expresses SHH medulloblastoma and neuronal differentiation markers (for example LMO4, FOXS1, HHIP), representing tumour cells with neuronal-like traits in highly vascular tumour regions. **b** Subclustering of reactive astrocyte-like / stress-response cells. Further subclustering of Cluster 0 identifies four transcriptionally and spatially distinct states. Subcluster 2 expresses reactive glial markers (*TNR, GFAP, PTPRZ1, BCAN, C1orf61*), consistent with reactive gliosis in brain parenchyma adjacent to tumour. Subcluster 1 co-expresses neuronal differentiation and proliferation markers (*INSM1, NEUROD1, NHLH1, STMN2, CCND2*), indicating a mixed differentiating/proliferative phenotype. Subcluster 3 is enriched for mesenchymal-like and vascular ECM genes (*VIM, LGALS1, COL4A1, COL4A2*), suggesting tissue damage and vascular involvement. Subcluster 0 shows high expression of ECM-remodelling and vascular markers (*MGP, COL12A1, LTBP2*), consistent with active stromal remodelling and angiogenesis. **c** Subclustering of extracellular matrix and stromal regions. Subclustering of ECM-rich and stromal areas delineates two principal states. Subcluster 0 is enriched for ECM and angiogenesis markers (*DCN, COL3A1, COL1A1, COL1A2*), reflecting tumour–stroma interactions and vascular remodelling. Subcluster 1 expresses neuronal-like and differentiated tumour markers (*INSM1, SOX4, RELN, SEZ6L, FGF14, TUBB2B*), indicating pockets of differentiated tumour cells embedded within the stromal microenvironment.

**Supplementary Fig. S9.**
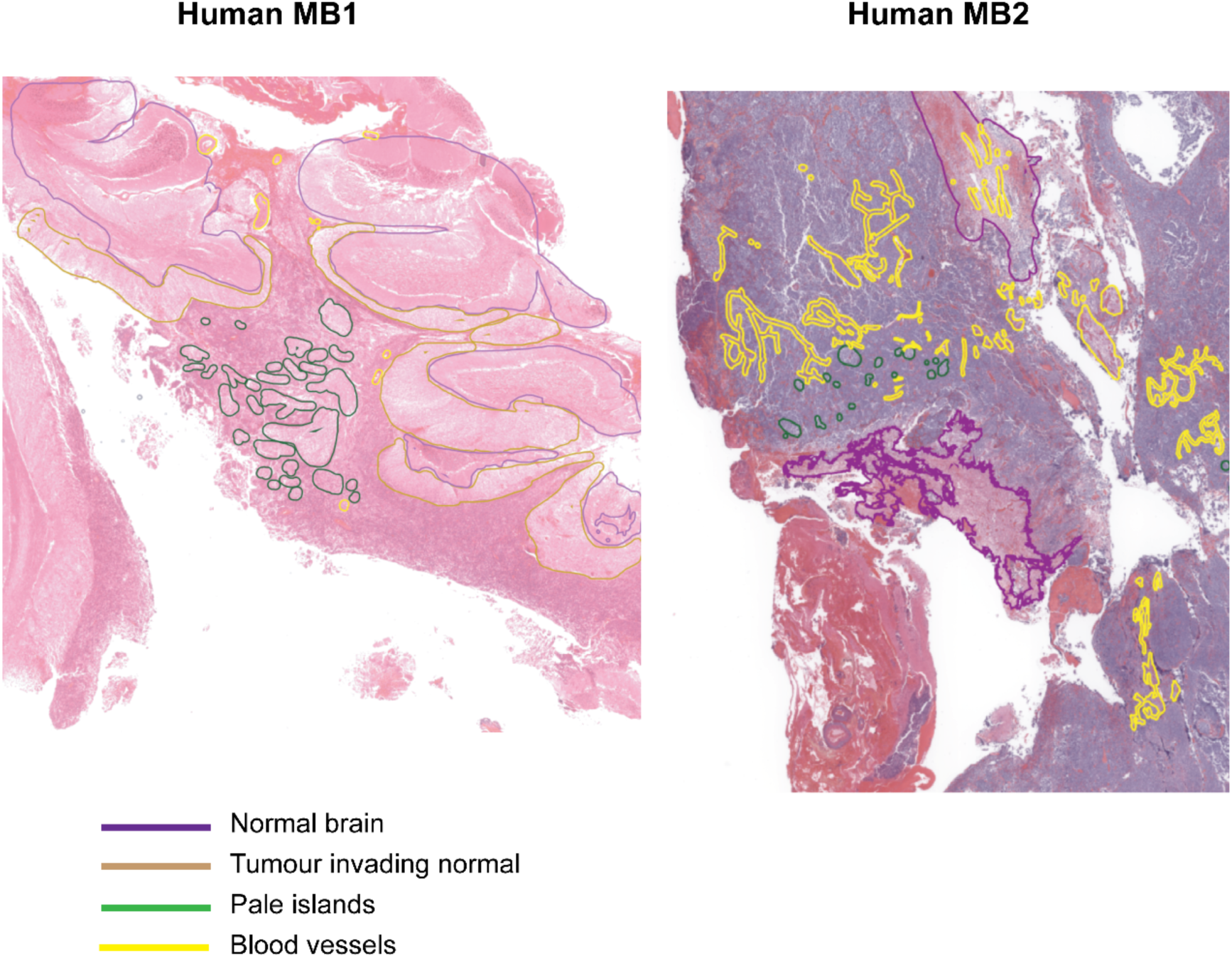
Histopathological annotation of human SHH medulloblastoma specimens. H&E-stained sections from two human SHH medulloblastomas (Human MB1 and Human MB2) with manual annotations by a neuropathologist. Overlays delineate normal brain parenchyma (purple), tumour invading normal brain (dark green), pale islands (light green) and blood vessels (yellow), providing structural context for spatial transcriptomic and metabolomic analyses in these samples.

**Supplementary Fig. S10.**
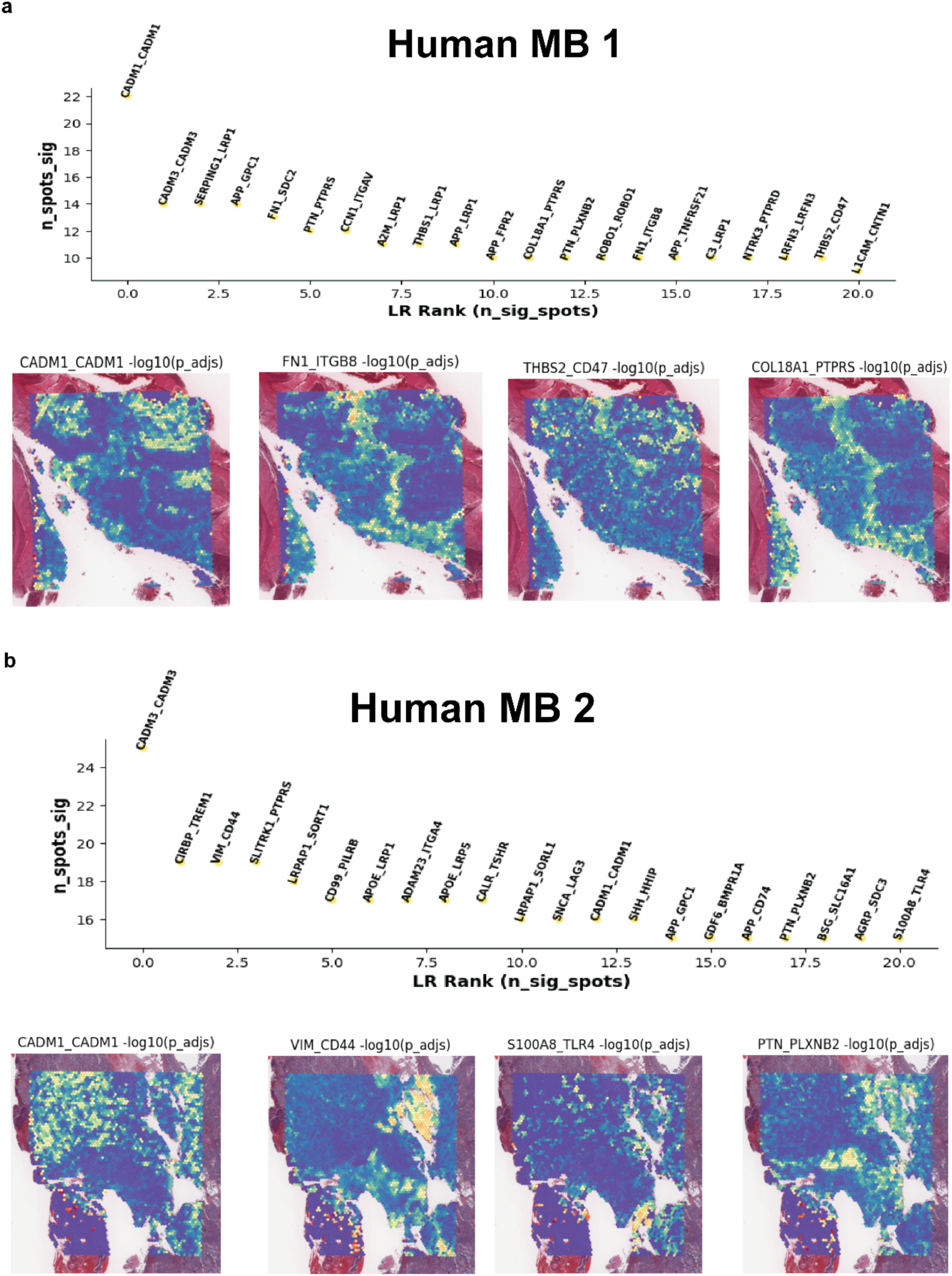
Spatial ligand–receptor signalling in human SHH medulloblastoma. **a** Human MB1. Top, ranked ligand–receptor (LR) interactions inferred from spatial transcriptomics in Human MB1, ordered by the number of significant spots and coloured by statistical significance (–log₁₀ adjusted P). Bottom, spatial feature plots for selected LR pairs (for example *CADM1–CADM1, F11R–ITGB8, THBS2–CD47, COL18A1–PTPRS*), showing localisation of high LR activity (yellow) within the tumour and at tumour–brain and perivascular interfaces. **b** Human MB2. Top, ranked LR interactions in Human MB2, displayed as in a. Bottom, spatial maps of representative LR pairs enriched in this specimen (for example *CADM1–CADM1, VIM–CD44, S100A4–TUBB4, PTN–PLXNB2*), illustrating recurrent tumour–stromal communication hubs across the section. Together, these analyses highlight conserved and sample-specific ligand–receptor networks potentially mediating tumour invasion and microenvironmental crosstalk.

**Supplementary Fig. S11.**
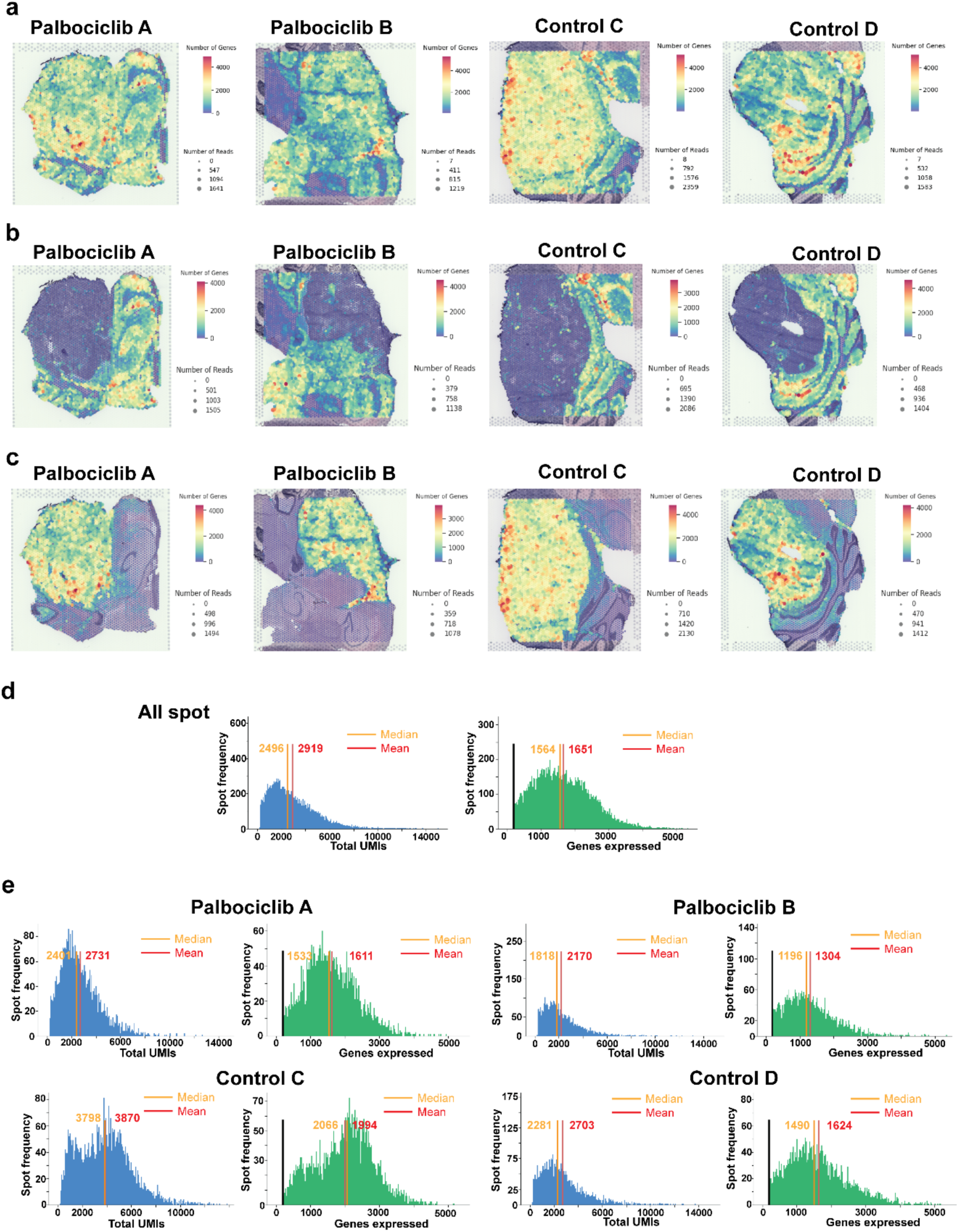
Quality assessment of Visium spatial transcriptomics in MB-PDOX brains. **a** Spatial maps of total genes detected per Visium spot in Palbociclib-treated (Palbociclib A, Palbociclib B) and control (Control C, Control D) MB-PDOX brains. Spots are coloured by gene count, illustrating overall sequencing depth and spatial heterogeneity in transcript capture. **b** Spatial distribution of human transcripts, showing that human gene expression is confined to the tumour regions in each section. **c** Spatial distribution of mouse transcripts from the same sections, highlighting surrounding mouse brain and stromal compartments. **d** Global quality- control histograms for all spots across samples, showing the distribution of total UMIs (left, blue) and number of detected genes (right, green). Red and orange lines indicate mean and median values, respectively; a minimum threshold of 200 genes per spot (black line) is used to exclude low-quality spots. **e** Sample-wise histograms of total UMIs (blue) and detected genes (green) for each MB-PDOX brain, with mean and median values indicated, confirming consistent sequencing depth and gene detection across treated and control datasets. Across the four MB-PDOX brains, Visium profiling yielded transcriptomes from 14,982 barcoded array spots. Gene counts per spot were consistently high (median 1,196–2,731), indicating robust RNA preservation and capture across sections.

## Data availability

## Code availability

The code can be found at: https://github.com/BiomedicalMachineLearning/SpatialMetabolomics https://github.com/GenomicsMachineLearning/SpaMTP

## Acknowledgements

The authors would like to thank the patients and families who contributed brain tumour tissue. This research was carried out in part at the Translational Research Institute, Woolloongabba, QLD, 4102, Australia. We thank to Queensland pathology for providing the samples and service for our study. Samples were sequenced by Institute for Molecular Bioscience sequencing facility. We thank the participants of this facility and acknowledge the contributions of the investigators to this study. We thank the technical supports from the QIMRB National Centre for Spatial Tissue and AI Research (NCSTAR) and the ACRF Centre for Optimised Cancer Therapies.

## Author information

These authors contributed equally: Tuan Vo and Cedric S. Cui

## Contributions

Q.N., J.L., T.V., C.S.C., J.F., L.G. designed the research project and research methods; Y.D., A.M.R.J. prepared the PDOX model; T.V., N.L. performed cryo-sectioning; C.S.C., B.H. performed MALDI-MSI; T.V., H.V. performed spatial transcriptomics and sequencing; A.X., T.V. performed Xenium and H&E staining; ; T.R. provided samples annotation and biology; M.H. provided expertise in mass spectrometry; T.V, C.S.C., A.T., E.J.K., H.V., Y.C, A.C., X.T., Q.N. contributed new datasets, reagents and analytical tools; T.V., A.T., Y.C., A.C., X.T. analysed data; T.V., C.S.C., A.T., Y.C., X.T., J.F., J.L., Q.N. wrote the manuscript. M.H., T.R., B.W. edited the manuscript; All authors read and approved the final manuscript. Q.N. and J.L. conceived the study and co-supervised the work.

## Ethics declarations

All animal procedures were approved by The University of Queensland Molecular Biosciences Animal Ethics Committee (IMB/386/18) and conducted in accordance with institutional guidelines. Human paediatric medulloblastoma FFPE tissue samples (n = 5) were obtained as de-identified archival specimens from Pathology Queensland, Central Laboratory (Herston, QLD, Australia) under approval from The University of Queensland Human Research Ethics Committee (2015001410). All procedures involving human tissue were performed in accordance with institutional guidelines and the tenets of the Declaration of Helsinki. Additional human transcriptomic data used in this study were obtained from previously published, publicly available datasets.

## Peer review

**Additional information**

**Supplementary information**

